# N-terminal proteoforms may engage in different protein complexes

**DOI:** 10.1101/2023.01.17.524352

**Authors:** Annelies Bogaert, Daria Fijalkowska, An Staes, Tessa Van de Steene, Marnik Vuylsteke, Charlotte Stadler, Sven Eyckerman, Kerstin Spirohn, Tong Hao, Michael A. Calderwood, Kris Gevaert

## Abstract

Alternative translation initiation and alternative splicing may give rise to N-terminal proteoforms, proteins that differ at their N-terminus compared to their canonical counterparts. Such proteoforms can have altered localizations, stabilities and functions. While proteoforms generated from splice variants can be engaged in different protein complexes, it remained to be studied to what extent this applies to N-terminal proteoforms. To address this, we mapped the interactomes of several pairs of N-terminal proteoforms and their canonical counterparts. First, we generated a catalogue of N-terminal proteoforms found in the HEK293T cellular cytosol from which 22 pairs were selected for interactome profiling. Additionally, we provide evidence for the expression of several N-terminal proteoforms, identified in our catalogue, across different human tissues as well as tissue-specific expression, highlighting their biological relevance. Protein-protein interaction profiling revealed that the overlap of the interactomes for both proteoforms is generally high, showing their functional relation. We also showed that N-terminal proteoforms can be engaged in new interactions and/or lose several interactions compared to their canonical counterpart, thus further expanding the functional diversity of proteomes.

## Introduction

Eukaryotic protein-coding genes give rise to several protein variants, or proteoforms, through various mechanisms including genetic alterations, alternative promotor usage during transcription, alternative splicing during mRNA maturation, use of alternative initiation codons and stop codon read-through during translation, and numerous co-and post-translational modifications [1–3]. Crosstalk between these mechanisms greatly expands a proteome’s complexity [4]. Studies in our lab revealed that 10-20% of protein N-termini in several human and mouse cells point to alternative translation initiation and/or alternative splicing [5]. Such N-terminal (Nt-) proteoforms thus stem from the same gene but differ at their N-terminus. The majority of Nt-proteoforms are truncated at the N-terminus relative to the canonical form however, up to 6% have extended N-terminal regions presumably caused by ribosomes starting translation from codons in the annotated 5’UTR. Nt-proteoforms can also carry modified N-termini different from those of the canonical protein [2, 6, 7].

Nt-proteoforms have long been overlooked and studies on their biological function are just now emerging [2, 8–11]. Nt-proteoforms may have different functions as the N-terminus of a protein steers several protein features such as half-life and protein localization [12, 13]. Concerning the latter, many targeting signals reside at a protein’s N-terminus and, consequently, N-terminally truncated or extended proteoforms may lose or gain targeting signals, causing such proteoforms to reside at different subcellular localizations [8, 14–22]. Several Nt-proteoforms with such altered subcellular localization are iso-functional, but thus active in different compartments [16, 17]. Our lab and the Kuster lab showed that pairs of Nt-proteoforms originating from the same gene can possess different stabilities in cells [9, 23, 24]. Of note, mounting evidence indicates that alternative translation initiation is regulated in response to a variety of stress stimuli and/or in a tissue and a cell developmental specific manner [3, 25, 26]. Van Damme *et al*. (2014) also showed that alternative translation initiation sites are generally conserved among eukaryotes, hinting to their possible biological impact [5]. In addition, several Nt-proteoforms have already been linked to human diseases, illustrating their potential for therapeutic intervention, diagnosing and prognosing disease [2, 27]. Other studies showed that N-terminal proteoforms may have altered functionalities [11, 25, 26, 28-33]. An example is the regulator of G-protein signaling (RGS2) which was reported to give rise to four different N-terminal proteoforms starting at methionines 1, 5, 16 or 33. The proteoforms starting at positions 16 or 33 have an impaired inhibitory effect on type V adenylyl cyclase (ACV) compared to the full-length protein and it was suggested that these N-terminal RGS2 proteoforms are part of a novel negative feedback control pathway for adenylyl cyclase signaling [29]. Different studies illustrated that Nt-proteoforms can interact with proteins other than the interaction partners of their canonical protein. For example, the fibroblast growth factor-2 (FGF-2) exists in multiple proteoforms: a low molecular weight Nt-proteoform (18 kDa) generated upon alternative usage of a start codon, and at least two higher molecular weight proteoforms (21 and 23 kDa) generated upon translation starting from CUG codons located in the 5’ UTR of the corresponding transcript. The 18 kDa and 23 kDa proteoforms have different localizations and different functionalities. Moreover, the 23 kDa FGF-2 proteoform co-immunoprecipitated with the survival of motor neuron protein (SMN), whereas the 18 kDa proteoform did not. The authors hence concluded that SMN specifically interacts with the 23-kDa FGF-2 proteoform by binding to its Nt-extension [11].

Protein-protein interactions (PPIs), either stable or transient, are important for cellular functions and regulate cellular signaling [34]. Mapping of protein-protein interaction networks is thus essential to understand cellular processes and signaling pathways, as well as for defining the origin of several human diseases [35]. Several efforts were made to create huge databases that contain experimentally determined PPIs of different organisms, such as BioGrid [36] and STRING [37, 38]. As mentioned by Ghadie *et al.*, these databases typically assume that one gene encodes for one protein and ignore the effects of protein modifications, alternative splicing, alternative translation initiation and other mechanisms leading to proteoforms [39]. Recently, a global study revealed a huge impact of protein isoforms originating from alternative splicing on the composition of protein complexes [40] and this often in a tissue-specific way as most proteoforms are expressed in specific tissues and play a role in network organization, function and cross-tissue dynamics [39–41]. Hence, proteoforms may not be overlooked when studying protein-protein interactions.

Based on different reports focusing on single pairs of proteoforms, we hypothesized that different Nt-proteoforms can be engaged in different protein complexes. As indicated in a recent review [42], no systems-wide information about the interplay between specific proteoforms and protein complex formation is yet available, and as proteins mainly function as part of protein complexes, it is interesting to explore to what extent proteoforms indeed affect interactomes [42]. Our prime objective was to assess our hypothesis at a larger scale using a contemporary approach for characterizing protein complexes and, by unraveling the PPIs of Nt-proteoforms, we aimed to learn more about their functions and how they contribute to the global functional complexity of the proteome. Here, we first applied N-terminal COFRADIC [43] on the cytosol of HEK293T cells to construct a comprehensive catalogue of N-terminal cytosolic proteoforms. We then applied stringent filtering to select proteoforms pairs for interactome analysis by Virotrap (see **Figure 1**), a method to study protein-protein interactions that avoids cell lysis by exploiting the characteristics of the HIV-1 p55 GAG protein which leads to the production of virus-like particles (VLPs). Of note, Virotrap was shown to be a sensitive PPI method, as VLPs encapsulate and preserve the protein complexes, allowing the detection of weak and transient protein-protein interactions [44].

**Figure 1:**
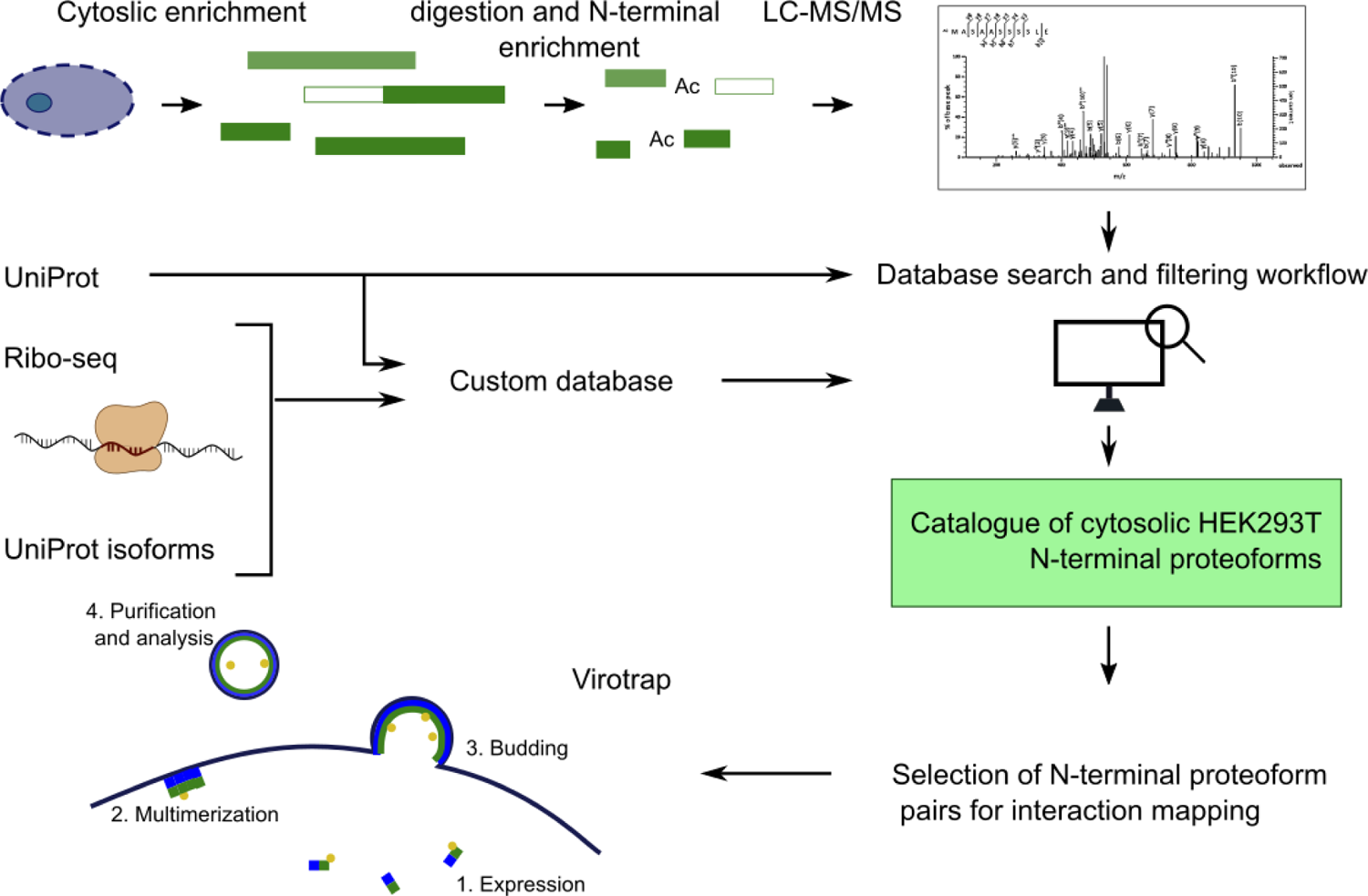
Overview of the experimental approach. HEK293T cells are grown followed by digitonin lysis to enrich for cytosolic proteins. Proteins are subsequently digested (in parallel using different proteases) and protein N-terminal peptides are enriched by COFRADIC, analyzed by LC-MS/MS and searched with two different databases: the UniProt database and a custom build database also including UniProt isoforms and Ribo-seq data. The data are then thoroughly filtered to generate a catalogue of N-terminal proteoforms and from this, pairs of canonical protein and N-terminal proteoforms are selected for interactome analysis by Virotrap.

In this study, we identified 3,306 protein N-termini in the cytosol of HEK293T cells of which 1,044 originate from N-terminal proteoforms, highlighting the prevalence of both alternative translation initiation and alternative splicing. We provide evidence for the existence of several of these N-terminal proteoforms in other cells and tissues, supporting their biological relevance. Virotrap-based interactome analysis of 20 carefully selected pairs of N-terminal proteoforms and their canonical protein revealed that N-terminal proteoforms share the majority of their interactions with the canonical protein, yet also have their own set of unique interaction partners, while other interactions can be lost.

## Materials and Methods

See Supplementary Materials and Methods for additional information.

### Cell culture

Human embryotic kidney cells (HEK293T cells) were cultured at 37 °C and 8% CO_2_ in DMEM medium supplemented with 10% fetal bovine serum (unless specified otherwise).

### Generation of a catalogue of N-terminal proteoforms in HEK293T cells

All experimental details on the sample preparation and LC-MS/MS analysis can be found in Supplementary Materials and Methods. From the set of identified peptides, N-terminal peptides were selected and classified as described in [45]. Different here is that data from two database searches are combined. The merging of the data from the two searches was done per protease after accession sorting, which ensures that peptides reported by different searches (or proteases) are matched to the same accession, allowing a straightforward merge. During this merge, all information was retained (e.g. all listed accessions are retained in the isoform column) and, after merging, the workflow was followed as outlined in [45] to filter for N-terminal peptides stemming from translation events.

### Generation of a cytosolic proteome map of HEK293T cells

Cytosolic extracts (in triplicate) were prepared from 10^7^ HEK293T cells similar as described in Supplementary Materials and Methods. The protein concentration was measured using a Bradford assay (Bio-Rad) and 500 µg of protein material was used to prepare samples for mass spectrometry analysis. Proteins were digested with endoproteinase-LysC (Promega, 1:100, w:w) for 4 h at 37 °C and then with trypsin (Promega, 1:100, w:w) overnight at 37 °C. Samples were acidified to a pH<3 by adding TFA to a final concentration of 1% (v/v). Following a 15-minutes incubation on ice, the samples were centrifuged for 15 min at 1750 xg at room temperature. The peptides-containing supernatant was further purified using SampliQ C18 100 mg columns (Agilent) according to the manufacturer’s instructions. Purified peptides were dried completely by vacuum drying, re-dissolved in 200 µl loading solvent A (0.1% TFA in water/acetonitrile (98:2, v/v)). The peptide concentration was measured on a Lunatic microfluidic device (Unchained Labs) and from each replicate, 50 µg of peptide material was injected for fractionation by RP-HPLC (Agilent series 1200) connected to a Probot fractionator (LC Packings). Peptides were first loaded in solvent A on a 4 cm pre-column (made in-house, 250 µm internal diameter (ID), 5 µm C18 beads, Dr. Maisch) for 10 min at 25 µl/min and then separated on a 15 cm analytical column (made in-house, 250 µm ID, 3 µm C18 beads, Dr. Maisch). Elution was done using a linear gradient from 100% RP-HPLC solvent A (10 mM ammonium acetate (pH 5.5) in water/ACN (98:2, v/v)) to 100% RP-HPLC solvent B (70% ACN, 10 mM ammonium acetate (pH 5.5)) in 100 min at a constant flow rate of 3 µL/min. Fractions were collected every minute between 20 and 85 min and pooled to generate a total of 10 samples for LC-MS/MS analysis. All 10 fractions were dried under vacuum in HPLC inserts and stored at −20 °C until use.

Peptides were re-dissolved in 20 µl of 0.1% TFA and 2% ACN and 15 µl was injected for LC-MS/MS analysis on an Ultimate 3000 RSLC nano LC (Thermo Fisher Scientific, Bremen, Germany) in-line connected to a Q Exactive mass spectrometer (Thermo Fisher Scientific). The peptides were first loaded on a trapping column (made in-house, 100 μm internal diameter (I.D.) × 20 mm, 5 μm beads C18 Reprosil-HD, Dr. Maisch) with loading solvent (0.1% TFA in water/ acetonitrile, 2/98 (v/v)). After 4 min, a valve switch put the loading column in-line with the analytical pump to load the peptides on a 50 cm µPAC™ column with C18-endcapped functionality (PharmaFluidics) kept at a constant temperature of 35 °C. Peptides were separated with a non-linear gradient from 98% solvent A’ (0.1% formic acid in water) to 30% solvent B′ (0.1% formic acid in water/acetonitrile, 20/80 (v/v)) in 70 min, further increasing to 50% solvent B’ in 15 min before reaching 99% solvent B’ in another 1 min. The column was then washed at 99% solvent B’ for 5 min and equilibrated for 15 min with 98% solvent A’. The flow rate was kept constant at 300 nl/min for the entire run, except for the first 9 min during which the flow rate was set to 750 nl/min.

The mass spectrometer was operated in data-dependent, positive ionization mode, automatically switching between MS and MS/MS acquisition for the five most abundant peaks in a given MS spectrum. The source voltage was 2.6 kV, and the capillary temperature was 275 °C. One MS1 scan (m/z 400−2,000, AGC target 3 × 10^6^ ions, maximum ion injection time 80 ms), acquired at a resolution of 70,000 (at 200 m/z), was followed by up to 5 tandem MS scans (resolution 17,500 at 200 m/z, AGC target 5 × 10^4^ ions, maximum ion injection time 80 ms, isolation window 2 m/z, fixed first mass 140 m/z, spectrum data type: centroid) of the most intense ions fulfilling predefined selection criteria (intensity threshold 1.3xE^4^, exclusion of unassigned, 1, 5-8, >8 positively charged precursors, peptide match preferred, exclude isotopes on, dynamic exclusion time 12 s). The HCD collision energy was set to 25% Normalized Collision Energy and the polydimethylcyclosiloxane background ion at 445.120025 Da was used for internal calibration (lock mass).

The generated MS/MS spectra were processed with MaxQuant (version 1.6.1.3) using the Andromeda search engine with default search settings, including a false discovery rate set at 1% on both the peptide and protein level. Spectra were searched against the sequences of the human proteins in the Swiss-Prot database (release Jan 2019). Matching between runs was enabled and performed on the level of pre-fractionated peptides (fraction 1-10) such that it was enabled for each preceding and each following peptide fraction. The enzyme specificity was set at trypsin/P, allowing for two missed cleavages. Variable modifications were set to oxidation of methionine residues and N-terminal protein acetylation. Carbamidomethylation of cysteine residues was put as a fixed modification. All other settings were kept standard. Proteins were quantified by the MaxLFQ algorithm integrated in the MaxQuant software [46]. A minimum of two ratio counts and both unique and razor peptides were considered for protein quantification, leading to the identification of 3,933 proteins across all samples.

Further data analysis was performed with the Perseus software [47] (version 1.6.3.4) after uploading the protein groups file from MaxQuant. Proteins only identified by site and reverse database hits were removed as well as potential contaminants. To be able to perform GO (Gene Ontology), KEGG, Pfam, Corum and Keyword term enrichment, available annotations were uploaded from the *Homo sapiens* database (release date 2018-04-06). The replicate samples were grouped and the LFQ intensities were log_2_ transformed. Proteins with less than two valid values in at least one group were removed and missing values were imputed from a normal distribution around the detection limit (with 0.3 spread and 1.8 down-shift). This led to the identification of 3,045 proteins. The efficiency of the cytosol enrichment was evaluated by loading all identified UniProt identifications in the retrieveID/mapping tool on their website and checking the amount of proteins listed with the GOCC term cytosol.

### Selection of N-terminal proteoform pairs for interactome analysis

Nt-peptides selected using the KNIME workflow outlined in [43] and Supplementary Materials and Methods were further annotated and curated in R version 4.1.0.

Part I – annotation of genomic features. Human gene and transcript annotations were downloaded from Ensembl BioMart Archive Release 98 (September 2019) and proteoform accessions were linked to gene identifiers, names, biotypes and transcript support levels (TSL). Subsequently, Nt-peptides were mapped to genomic positions and a peptide BED file was created. For this purpose, Nt-peptides matching a Ribo-seq-derived protein accession were represented as transcript coordinates and converting to genomic coordinates using Proteoformer output SQL database (for translation initiation site (TIS) distance to transcript start), R packages biomaRt (2.50.3), GenomicFeatures (1.46.5), rtracklayer (1.54.0) and RMariaDB (1.2.2). Genomic coordinates were generated for UniProt-mapped peptides only if the entire UniProt proteoform was identical to the Ensembl annotated proteoform. Therefore, 109 peptides could not be mapped to the genome. Alternative translation initiation events (>1 TIS in the same exon) were distinguished from otherwise possible alternative splicing (TISs in different exons) by extracting the exon ranks of each Nt-proteoform start site and the corresponding aTIS on the same transcript (if available). Furthermore, we report the exact Nt-proteoform start codon, frame and distance to the corresponding aTIS.

Part II – annotation and prediction of protein sequence features. UniProt annotations of human canonical proteins were downloaded in January 2020. Positions of the following features: signal peptide, transit peptide, propeptide, region, motif, coiled coil, compositional bias, repeat and zinc finger were converted to protein sequence ranges using GenomicRanges (version 1.46.1). To determine sequence features annotated in UniProt that Nt-proteoforms had lost, Nt-peptides were exactly matched to canonical human UniProt proteins using dbtoolkit (version 4.2.5) [48]. For Nt-peptides that mapped internally to a canonical proteoform, we determined the lost (Nt-truncated) region. For 5’UTR-extended proteoforms (compared to the canonical proteoform), we considered any sequence feature spanning position 1 or 2 in the canonical protein to be no longer N-terminal in the 5’ extended proteoform and thus lost. Otherwise, for Nt-peptides not exactly mapping to any UniProt protein, alignment of the entire Nt-proteoform to one UniProt reference sequence was performed (see Table S4 column “proteoform.sequence” and “sequence.feature.reference.ID”, respectively) using Biostrings version 2.62.0 pairwiseAlignment function with the following parameters: substitutionMatrix = BLOSUM62, gapOpening = −9.5, gapExtension =-0.5, scoreOnly = FALSE, type=“overlap”. Selection criteria of the reference sequence for alignment were as follows: UniProt isofoms were aligned to the matching canonical accession. Ensembl proteoforms were aligned to the UniProt protein from the same gene. In absence of any reliable UniProt reference (e.g. Ensembl NTR proteoforms), no alignment was performed. From these alignments, sequence ranges that were lost compared to the UniProt reference protein were extracted. We overlapped such lost regions (from exact matching or alignment) with sequence features in the UniProt database (see Table S4 column “lost.UniProtDB.sequence.feature”). We also determined if Nt-truncated proteoforms could be derived from N-terminal processing (removed signal, transit or pro-peptide) considering a ± one amino acid margin of error (see Table S4 column “signal.transit.propeptide.processing_distance”). For Nt-peptides without an exact match to an UniProt protein, we scanned for short linear motifs in the proteoform and their selected UniProt reference using Eukaryotic Linear Motif (ELM) resource API [49]. From the alignments described above, we extracted sequence stretches that are lost compared to the UniProt reference proteins or sequence stretches gained by the proteoform, and reported the ELMs predicted in these regions (see Table S4 columns “lost_ELM”and “gained_ELM”, respectively). TopFIND [50] was used to determine if the Nt-peptides could have been derived from post-translational processing (thus leaving out co-translational processing by methionine aminopeptidases; TopFIND results are presented in Table S4 column “Cleaving.proteases” and “Distance.to.last.transmembrane.domain.shed”). Additionally, Nt-proteoforms that could derive from N-terminal dipeptidase cleavage are marked; see Table S4 column “dipeptidase”. Finally, to fully explore sequence similarity of Nt-proteoforms without an exact UniProt match to other known or predicted proteins in the NCBI database, we performed a BLAST analysis against the human UniProt or the human non-redundant proteins (NCBI, July 2020). Hits with over 80% of protein sequence identity over 50% of the length are reported (see Table S4 columns “blast.vs.uniprot”, “blast.vs.nonredundant”). The molecular weights of Nt-proteoforms and their annotated counterparts were calculated using R package Peptides version 2.4.4.

Part III – survey of complementary data(bases). Curated, experimentally determined protein interactions were downloaded from BioGRID version 3.5.182 [51], matched by gene name and reported in Table S4 column “biogrid”. Human genetic phenotypes and disorders available from OMIM [52] were matched by gene identifier using biomaRt version 2.50.3, release 98 (see Table S4 column “omim”). Prior experimental evidence for Nt-proteoform expression in human (primary) cells reported by Van Damme *et al.* 2014 was included for matching N-termini (allowing for Met processing, see Table S4 column “VanDamme.2014”) [5]. Cytosolic expression associated with the matching gene was reported in Table S4 column “cytosolic.in.HEK.proteome” when confirmed by three independent sources: our own cytosolic proteomics data (see above), gene ontology GOSlim annotation of cellular component (containing “cytosol” and “cytoplasm”, whilst excluding “organelle”) [53] and cytosolic subcellular localization determined by immunostaining from The Human Protein Atlas [54].

Genes with multiple proteoforms were classified into four categories: 1. annotated + alternative TIS, 2. multiple alternative TIS, 3. multiple annotated TIS and 4. one TIS, where canonical UniProt and Ensembl aTIS were considered annotated TIS. Subsequently, we calculated a TIS score for each alternative Nt-proteoform. High confident TIS (according to our KNIME workflow), > 50% acetylation, spectral count > 1, several Nt-peptides pointing to the same Nt-proteoform, database or Ribo-seq evidence for TIS, identification in multiple protease conditions (trypsin, chymotrypsin or endoproteinase GluC), non-AUG start codon in 5’UTR, lost or gained ELM, presence in the Van Damme *et al.* 2014 dataset, genetic disease from OMIM or no interactions known in BioGRID all increased the score by 1. Truncation of less than 50 % of protein length and a UniProt domain lost additionally increased the score by 3. The score however dropped to 0 when we suspected protease cleavage (from TopFIND, through dipeptidase activity, signal, transit or pro-peptide processing) gave rise to the Nt-proteoform. A gene score was further calculated as the sum of Nt-proteoform scores of a given gene. Gene scores were kept only for genes of the following categories: 1. annotated + alternative TIS, 2. multiple alternative TIS and 4. one TIS, if we had orthogonal evidence of cytosolic localization, thus prioritizing genes with novel proteoforms. If the gene score was >1 it is included in the Supplementary Table S3 (Tab 2, gene score >0).

The list of 372 genes with a score >1 was trimmed to a manageable list for Virotrap analysis using prioritization and selection criteria. These criteria, ranked according to their importance, are the following. 1. High confident N-terminal proteoforms are prioritized over low confident proteoforms. The former are either assigned as high confident following our KNIME filtering strategy or for which there was extra evidence supporting their synthesis. Such evidence can either be a known UniProt isoform, extra Ribo-seq evidence supporting the TIS, its reporting in the dataset of Van Damme *et al.* [5], and its detection on Western blots as available in HPA (see next paragraph for details). 2. Proteoforms with a loss or gain of a known domain or eukaryotic linear motif (ELM) were given higher priority. 3. Proteoforms with a truncation or extension of >20 amino acids or >50 amino acids for long proteins (> 700 amino acids) were preferred over shorter truncations. 4. The more evidence for the cytosolic localization of the proteoform, the higher its priority as Virotrap is restricted to cytosolic proteins. To increase the overall confidence for the cytosolic localization of a proteoform, we also relied on our own cytosolic proteome map. 5. Non-structural proteins such as enzymes were prioritized over structural proteins (e.g., cytoskeletal proteins) as Virotrap encountered difficulties using structural proteins as baits. 6. Proteins that have a known association with a human disease according to the OMIN database were prioritized over proteins without such a link. 7. Proteoforms resulting from translation out-of-frame from the start site of the canonical protein sequence were considered as novel proteins, thus not as Nt-proteoforms, and were not selected for further analysis.

To further evaluate the identified N-terminal proteoforms, we used Western blot (WB) data available in The Human Protein Atlas (HPA, https://www.proteinatlas.org/). From the 372 genes with a score > 1, we evaluated if the N-terminal proteoform could be detectable on WB (based on the actual mass difference and the likelihood that the epitope recognized by the antibody is still present in the proteoform). For 138 out of the 372 genes, the N-terminal proteoforms would potentially be detectable and these genes were further evaluated here. Information on all antibodies tested for the protein products of these genes (also including unpublished antibodies) was retrieved from the HPA LIMS. A WB score was retrieved. This score ranges between 1-7 and points to the following: 1) Single band corresponding to the predicted size (±20%). 2) Band of predicted size (±20%) with additional bands present. 3) Single band larger than predicted size (+20%), but partly supported by experimental and/or bioinformatics data. 4) No bands detected. 5) Single band differing more than ±20% from predicted size and not supported by experimental and/or bioinformatics data. 6) Weak band of predicted size but with additional bands of higher intensity also present and 7) only bands that do not correspond to the predicted size (extra information can be found here: https://www.proteinatlas.org/learn/method/western+blot). Genes yielding protein products with a WB score of two, for at least one of the antibodies raised again these protein products, were further checked. Although scores 6 and 7 also indicate extra bands (and thus potential Nt-proteoforms), the confidence in these results is lower and these were thus not considered further. For the genes giving rise to protein products with a score of two, the Western blots generated were checked on The Human Protein Atlas website (www.proteinatlas.org) for a protein band with a size corresponding to the N-terminal proteoform identified. When such bands were found, this further increased the overall confidence score for this N-terminal proteoform and was thus considered in the final selection.

The prioritization/selection and HPA information reduced the list of genes to 85 (pairs of canonical proteins and one or two proteoforms) and from this list, 22 genes were selected for Virotrap analysis that were all top-ranked as they met most to all selection criteria and some also contained HPA WB evidence.

### Tissue expression of Nt-proteoforms evaluated through re-analysis of public proteomics data

Mass spectrometry data of the draft human proteome map developed by the Pandey group [55], composed of 30 histologically normal human samples including 17 adult tissues, 7 fetal tissues and 6 purified primary hematopoietic cells, were downloaded from PRIDE project PXD000561 and searched with ionbot version 0.8.0 [56]. Of the 30 samples, each was processed by several sample preparation methods and MS acquisition pipelines to generate 84 technical replicates. We first generated target and decoy databases from our custom-build database containing UniProt canonical and isoform entries appended with Ribo-seq derived protein sequences [46]. Next, we searched the mass spectrometry data with semi-tryptic specificity, DeepLC retention time predictions [57] and protein inference enabled, precursor mass tolerance set to 10 ppm and a q-value filter of 0.01. Carbamidomethylation of cysteines was set as a fixed modification, oxidation of methionines and N-terminal acetylation were set as variable modifications and an open-modification search was disabled. Downstream analysis was performed in R version 4.1.0 using dplyr (1.0.9), Biostring (2.26.0), GenomicRanges (1.46.1) and biomaRt (2.50.3, release 98). To constrict the results to the first-ranked PSM per spectrum, we used ionbot.first.csv output and filtered out decoy hits, common contaminants that do not overlap with target FASTA and used PSM q-value ≤ 0.01. Due to the complexity of our custom protein database, most PSMs were associated with several protein accessions. We sorted accessions to prioritize UniProt canonical followed by UniProt isoforms, followed by Ribo-seq, higher peptide count (in the whole sample), start (smallest start position first) and accession (alphabetically).

These steps yielded a filtered PSM table. Subsequently, we sorted PSMs by N-terminal modification (to prioritize N-terminally acetylated peptidoforms) and highest PSM score. Sorted PSMs were grouped by matched peptide sequence yielding a unique peptide table. Peptides were grouped by sorted accession to generate a protein table, complemented with sample and protein metadata (such as gene and protein names, descriptions). Per sample and replicate, we obtained a unique peptide count, spectral count and NSAF (normalized spectral abundance factor) quantification. Differential expression analysis across all tissues was performed using limma (3.50.3) based on log2NSAF values only for proteoforms found in all replicates of at least one tissue (9,644/26,159 proteoforms). Importantly, for non-canonical proteoforms, only peptides that do not map to canonical UniProt proteins via dbtoolkit (version 4.2.5) [48] were considered for the quantification. We extracted pairwise contrasts adult vs. adult; fetal vs. fetal and adult vs. fetal of the same tissue, considering only significant differences in expression with a Benjamini-Hochberg adjusted p-value of 0.05. Boxplots were created using ggplot2 (3.3.6), whereas heatmaps of Nt-proteoform expression were generated using pheatmap (1.0.12). To determine the row clustering, we used log2NSAF values converted to binary data as input for MONothetic Analysis (cluster version 2.1.3).

### Generation of Virotrap clones

Gag-bait fusion constructs were generated as described [44]. The coding sequences for the full-length protein were either ordered from IDT (gBlocks gene fragments, as was the case for the following constructs: CAPRIN1, SPAST, PRUNE, SORBS3, FNTA, CAST, RARS, UBXN6, PAIP1, PXN and NTR protein) or generated cDNA was used as template for PCR amplification using AccuPrime^TM^ *pfx* DNA polymerase (Invitrogen) and ORF-specific primers. cDNA was generated by isolating RNA from 5×10^6^ HEK293T cells with the Nucleospin RNA isolation Mini kit (Macherey-Nagel)according to the manufacturer’s instructions. 500 ng of isolated RNA was then used as input for generating cDNA and cDNA synthesis was performed using the PrimeScript RT kit (Takara Bio) according to the manufacturer’s instructions. The generated PCR products for the full-length proteins were transferred into the pMET7-GAG-sp1-RAS plasmid by classic cloning with restriction enzymes (EcoRI and XbaI) or In-Fusion seamless cloning (Takara Bio) when the genes contained internal restriction sites for EcoRI and XbaI (which was the case for CSDE1, NTR and UBAC1).

The N-terminal proteoforms were amplified from the corresponding generated pMET7-GAG-sp1-FL plasmid of each gene using the AccuPrime^TM^ *pfx* DNA polymerase (Invitrogen) with proteoform-specific primers. However, some proteoforms also contain extensions or large internal deletions and these were ordered from IDT (gBlocks gene fragments, as was the case for the following constructs: the two N-terminal proteoforms of CAST (P20810-4 and P20810-8), N-terminal proteoform of PXN (P49023-4), an N-terminally extended proteoform of UBE2M and one of the two N-terminal proteoforms of SPAST (Q9UBP0-4)). The generated PCR products of the proteoforms were transferred into the pMET7-GAG-sp1-RAS plasmid as indicated above.

### Virotrap studies

For full details on the Virotrap protocol we refer to [44]. HEK293T cells were kept at low passage (<10) and cultured at 37 °C and 8% CO_2_ in DMEM, supplemented with 10% fetal bovine serum, 25 units/ml penicillin and 25 μg/ml streptomycin.

For Western blot validation of expression, 1.15×10^6^ HEK293T cells were seeded in 6-well plates the day before transfection. On the day of transfection, a DNA mixture was prepared containing the following: 0.82 μg bait construct (pMET7-GAG-sp1-bait), 0.046 μg pMD2.G and 0.093 μg pcDNA3-FLAG-VSV-G. In each experiment, eGFP and eDHFR were taken along as controls. For eGFP, normal expression amounts were used instead of maximal expression as used for the baits. For eDHFR, two expression amounts were used to allow comparison of bait intensities with those of the control. For normal expression, a DNA mixture was prepared containing the following: 0.48 μg control construct (either pMET7-GAG-sp1-eGFP or pMET7-GAG-sp1-eDHFR), 0.34 µg of pSVsport (mock vector), 0.046 μg pMD2.G and 0.046 μg pcDNA3-FLAG-VSV-G. Cells were transfected using polyethylenemine (PEI). After 6 h of transfection, the medium was refreshed with 2 ml of fresh growth medium. After 46 h, the cellular supernatant was collected and the cellular debris was removed from the harvested supernatant by 3 min centrifugation at 400 xg at room temperature. The cleared medium was then incubated with 10 μl Dynabeads MyOne Streptavidin T1 beads (Invitrogen) pre-loaded with 1 μg monoclonal Anti-FLAG BioM2-Biotin, Clone M2 (Sigma-Aldrich) according to the manufacturer’s protocol. After 2 h binding at 4 °C by end-over-end rotation, beads were washed twice with washing buffer (20 mM HEPES pH 7.5 and 150 mM NaCl) and the captured particles were released directly in 40 μl SDS–PAGE loading buffer. A 10-min incubation step at 65 °C before removal of the beads (by binding them to the magnet) ensured complete release and lysis of the VLPs.

Lysates of the producer cells were prepared by scraping the cells in 100 μl Gingras lysis buffer (50 mM HEPES-KOH pH 8.0, 100 mM KCl, 2 mM EDTA, 0.1% NP40 and 10% glycerol supplemented with 1 mM DTT, 0.5 mM PMSF, 50 mM Glycerophosphate, 10 mM NaF, 0.25 mM sodium orthovandate and Complete protease inhibitor cocktail (Roche)) after washing of the cells in chilled PBS. The lysates were cleared by centrifugation at 13,000 g, 4 °C for 15 min, to remove the insoluble fraction. The protein concentration was measured and 25 µg of protein material was mixed with SDS-PAGE loading buffer (diluted to a total volume of 30 µl). After heating to 95 °C for 5 min, both supernatant and lysate samples were loaded on Criterion XT 4-12% Bis-Tris gels (Biorad Laboratories). Each set of experiments also contained the GAG-EGFP expression control. Following SDS-PAGE, proteins were transferred to a PVDF membrane (Merck Millipore) after which the membrane was blocked using Odyssey Blocking buffer (LI-COR) diluted once with TBS-T (TBS supplemented with 0.1% Tween 20). Immunoblots were incubated overnight with primary antibodies against GAG (Abcam) and, for the lysates, also with primary antibody against actin (Sigma-Aldrich), serving as a loading control, in Odyssey Blocking buffer (PBS) diluted once with TBS-T. Blots were washed four times with TBS-T, incubated with fluorescently-labeled secondary antibodies, IRDye 800CW Goat Anti-Mouse IgG polyclonal 0.5 mg (LI-COR) and IRDye 680RD Goat Anti-Rabbit IgG polyclonal 0.5 mg (LI-COR) in Odyssey blocking buffer diluted once with TBS-T for 1 h. After three washes with TBS-T and an additional wash in TBS, immunoblots were imaged using the Odyssey infrared imaging system (LI-COR).

For LC-MS/MS analysis, all baits were divided into sets of maximally four baits and one control bait based on similar expression levels as judged from WB data. In total, we seven sets of baits were generated, and per set, always a FL and PR bait were included unless specified otherwise. The first set included eDHFR, CSDE1, MAVS and TSC22D3, the second set eDHFR, PAIP1, UBXN6 and ZFAND1, set number three consisted of eDHFR, SPAST (FL, PR1 and PR2) and UCHL1 (FL, PR1 and PR2), set number four eDHFR, CFL2 and FNTA, set number five eDHFR, AIMP1, PRPSAP1 and UBE2M, set six contained eDHFR, PRUNE1, PXN, UBAC1 and a protein from a non-translated region and the last set contained eDHFR, CACYBP, CAPRIN1 and EIF4A1. Each construct was analyzed in triplicate and for every replicate, the day prior to transfection, a 75 cm^2^ falcon was seeded with 9×10^6^ cells. Cells were transfected using polyethylenemine (PEI), with a DNA mixture containing 6.43 μg of bait plasmid (pMET7-GAG-bait), 0.71 μg of pcDNA3-FLAG-VSV-G plasmid and 0.36 μg of pMD2.G plasmid. Based on the WB results, we decided to use normal expression levels of eDHFR as their intensities were most similar to the intensities of the different baits. Thus, for the eDHFR control, cells were transfected with a DNA mixture containing 3.75 μg of eDHFR plasmid (pMET7-GAG-eDHFR), 2.68 µg of pSVsport plasmid, 0.71 μg of pcDNA3-FLAG-VSV-G plasmid and 0.36 μg of pMD2.G plasmid. The medium was refreshed after 6 h with 8 ml of DMEM supplemented with 10% FBS, 25 U/ml penicillin and 25 µg/ml streptomycin.

The cellular supernatant was harvested after 46 h and centrifuged for 3 min at 1,250 x g to remove debris. The cleared supernatant was then filtered using 0.45 μm filters (Merck Millipore). For every sample, 20 μl MyOne Streptavidin T1 beads in suspension (10 mg/ml, Thermo Fisher Scientific) were first washed with 300 μl wash buffer containing 20 mM Tris-HCl pH 7.5 and 150 mM NaCl, and subsequently pre-loaded with 2 μl biotinylated anti-FLAG antibody (BioM2, Sigma). This was done in 500 μl wash buffer and the mixture was incubated for 10 min at room temperature. Beads were added to the samples and the virus-like particles were allowed to bind for 2 h at room temperature by end-over-end rotation. Bead-particle complexes were washed once with 200 µl washing buffer (20 mM Tris-HCl pH 7.5 and 150 mM NaCl) and subsequently eluted with FLAG peptide (30 min at 37 °C; 200 μg/ml in washing buffer) and lysed by addition of Amphipol A8–35 (Anatrace) [58] to a final concertation of 1 mg/ml. After 10 min, the lysates were acidified (pH <3) by adding 2.5% formic acid (FA). Samples were centrifuged for 10 min at >20,000 x g to pellet the protein/Amphipol A8–35 complexes. The supernatant was removed and the pellet was resuspended in 20 µl 50 mM fresh triethylammonium bicarbonate (TEAB). Proteins were heated at 95 °C for 5 min, cooled on ice to room temperature for 5 min and digested overnight at 37 °C with 0.5 μg of sequencing-grade trypsin (Promega). Peptide mixtures were acidified to pH 3 with 1.5 µl 5% FA. Samples were centrifuged for 10 min at 20,000 x g and 7.5 µl of the supernatant was injected for LC-MS/MS analysis on an Ultimate 3000 RSLCnano system in-line connected to a Q Exactive HF Biopharma mass spectrometer (Thermo Scientific). Details on the LC-MS/MS settings and data-analysis can be found in Supplementary Materials and Methods.

The matrix containing all identified peptides (filtered on valid values and with imputed LFQ values) was exported from Perseus and statistical analysis was performed on GenStat (version V21, https://genstat21.kb.vsni.co.uk/). For each set, proteins were analyzed separately by fitting a linear model of the following form: response = µ + bait + error, where the response represents the log2-transformed LFQ intensity measured. The significance of the bait effect was assessed by a F-test and the significance of individual comparisons between the baits factor was assessed by a t-test. The performed pairwise comparisons are tabulated in **Supplementary Table S1**. Correction for multiple testing was done by estimating the false discovery rate (FDR) by modeling significance values as a 2-component mixture of Uniform and Beta or Gamma densities as implemented in Genstat v21.

Potential interaction partners of all baits (both full-length proteins and Nt-proteoform) were selected in pairwise contrasts between the bait and eDHFR control samples at an FDR of 0.01. The generated list of potential interaction partners was compared with known interaction partners listed in BioGRID [51], STRING [59] and IntAct [60]. Pairwise contrasts of interest between proteoforms were selected at an FDR of 0.05 and a difference of at least one log_2_ change. Further, when the difference in prey levels between the proteoform interactomes were at least two-fold, the difference required in order to be retained needed to be higher than the difference in levels between the bait proteoforms (on the side of the most intense proteoform). Only proteins also listed as candidate interaction partners in the comparisons of eDHFR control samples with the baits (FL and PR) were retained as potential differential interaction partners of the proteoforms.

### Generation of Y2H clones

Generation of Y2H clones was done as described [40]. All 22 selected pairs of full-length proteins and proteoforms were cloned into four different Y2H expression vectors by Gateway Cloning (Invitrogen). The full-length proteins and their N-terminal proteoforms were amplified from the corresponding pMET7-GAG-sp1-FL or pMET7-GAG-sp1-PR plasmids of each gene using the AccuPrimeTM pfx DNA polymerase (Invitrogen) and ORF-specific primers supplemented with *att*B sites. PCR products were transferred into pDONR221 by Gateway BP reaction (using BP clonase II enzyme mix, Invitrogen) according to the manufacturer’s instructions to generate entry clones. Entry clones were transformed and sequenced according to the manufacturer’s instructions. Then, all proteins were transferred from entry clones into each of the four destination vectors (pDEST-AD pDEST-AD-AR68, pDEST-AD-QZ213l and pDEST-DB, see [61] for vector details) by Gateway LR reaction (Gateway LR CLonase II Enzyme Mix, Invitrogen) according to the manufacturer’s instructions. The resulting expression vectors were used for Y2H.

### Y2H experiments

Y2H screening was performed as described [61]. All baits (coupled to the Gal4 DNA binding domain, DB) were tested against the hORFeome v9.1 collection of ∼17,408 ORF clones fused to the Gal4 activation domain (AD). Following first-pass screening, each bait was pairwise tested for interaction with the identified candidate partners. In this step, the interacting partners of any proteoforms of a gene were tested against all the proteoforms of that gene to eliminate false negatives due to sampling sensitivity. Pairs showing a positive result in the pairwise retest were PCR amplified and sequence confirmed (Sanger, Azenta) to confirm the identity of clones encoding each interacting protein.

### Generation of clones for AP-MS

Details can be found in Supplementary Materials and methods.

### Affinity-Purification Mass Spectrometry (AP-MS)

For AP-MS experiments, per experiment, 1.5 x 10^7^ HEK293T cells were seeded the day before transfection in a 150 mm dish. 7.5 μg of bait-FLAG DNA or 7.5 μg of eDHFR-FLAG DNA were transfected using polyethylenemine (PEI) as described above. 40 h post-transfection, cells were washed with ice-cold PBS and scraped in 1.5 ml of lysis buffer (containing 10 mM Tris-HCl pH 8, 150 mM NaCl, 1% NP-40, 10% glycerol, 1 mM sodium orhovanadate, 20 mM β-glycerophosphate, 1 mM NaF, 1 mM PMSF and cOmplete™ protease inhibitor cocktail). Cells were lysed on ice for 30 min and subsequently centrifuged 15 min at 4 °C at 20,000 x g. The supernatant was transferred to a new 1.5 ml protein LoBind Eppendorf tube and the protein concentration was measured using a Bradford assay. Sample volumes were adjusted with lysis buffer so that all samples were at the same protein concentration.

For every experiment, 10 μl MyOne Protein G beads (Invitrogen) were preloaded with 1 μg anti-FLAG (Sigma-Aldrich) and added to 350 µl of the cleared supernatant. Protein complexes were allowed to bind for 2 h by end-over-end rotation at 4 °C. The beads were washed twice with lysis buffer and three times with 20 mM Tris-HCl pH 8.0 and 2 mM CaCl_2_. Beads were re-suspended in 25 μl 20 mM Tris-HCl pH 8.0 and overnight incubated with 1 µg sequencing-grade modified trypsin at 37 °C. After removal of the beads, the samples were incubated for another 3 h with 250 ng trypsin. Samples were then acidified by 2% formic acid (f.c.) before LC-MS/MS analysis. Peptides were analyzed by LC-MS/MS using a Thermo Scientific Q Exactive HF operated similar as described for Virotrap (See Supplementary Materials and Methods). The generated MS/MS spectra were processed similar as described for Virotrap (see Supplementary Materials and Methods). Different here is that a newer version of MaxQuant was used (2.1.4.0) and that the database was supplemented with the sequence of eDHFR-FLAG. Overall, 2,705 proteins were identified over all samples. Based on the LC-MS/MS profiles, three samples were not further considered for analysis (eDHFR replicate C, PAIP1 FL replicate C and EIF4A1 FL replicate C).

Further data analysis was performed with the Perseus software [47] (version 1.6.15.0) after uploading the protein groups file from MaxQuant. Proteins only identified by site and reverse database hits were removed as well as potential contaminants. Replicate samples were grouped and proteins with less than three valid values in at least one group were removed, reducing the matrix to 1,903 protein identifications. Missing values were imputed using imputeLCMD (R package implemented in Perseus) with a truncated distribution with parameters estimated using quantile regression (QRILC). Imputed log2 LFQ values of AP-MS data were subjected to statistical analysis in R using limma version 3.50.3. Pairwise contrasts of interest between the different bait samples were retrieved at a significance level of alpha 0.05, corresponding to Benjamini–Hochberg adjusted *p* value (FDR) cutoff.

## Results

### 1. Construction of an N-terminal proteoform catalogue of the HEK293T cellular cytosol

To study the interactome of N-terminal proteoforms, we first constructed a catalogue of Nt-proteoforms of cytosolic proteins in HEK293T cells as Virotrap currently only functions in these cells and favors cytosolic proteins as baits. Such a decrease in proteome sample complexity also increases the possibility of identifying Nt-proteoforms [45]. Both Western blot and Gene Ontology Cellular Component data analysis indicated that we strongly enriched for cytosolic proteins (see **Supplementary Figure S1.A and B**). Given that Nt-proteoforms have different N-termini, we enriched for N-terminal peptides of the cytosolic proteins by N-terminal COFRADIC (omitting the SCX pre-enrichment step) [43]. In parallel, three different proteases, trypsin, chymotrypsin and or endoproteinase GluC, were used as this further increases the depth of analysis [62].

The LC-MS/MS data were searched using the UniProt database (restricted to human proteins) as well as a custom-build database which included all human UniProt proteins and UniProt isoforms, supplemented with protein sequences built from two HEK293T Ribo-seq datasets, and contains 103,020 non-redundant protein sequences [45]. As the custom database is significantly larger than the UniProt database (which only holds 20,356 proteins), and this negatively affects the FDR and the number of identified proteins [63, 64], we opted to combine the data of the two searches, thus boosting the total number of identifications. We evaluated how efficient N-terminal COFRADIC enriched for N-terminal peptides and found enrichments up to 60%, which is similar as previously reported [43] (see **Supplementary Figure S1.C**). Similar as reported in [45], the enrichment efficiency for chymotrypsin-digested samples is lower.

Further bioinformatics data curation and analysis, facilitating the selection of N-terminal proteoform pairs for interactome analysis, is summarized in **Figure 2A**. First, we applied stringent filtering on the identified peptides to retain N-terminal peptides originating from translation, as previously described [45] (see **Figure 2**, step 3 and its details in **Figure 2B**). We started by selecting N-termini pointing to database-annotated translation start sites and continued with inspecting N-terminal peptides starting upstream or downstream of annotated start sites to identify N-terminal proteoforms. Evaluation of candidate alternative N-termini favors peptides with co-translational Nt-acetylation, also considering its interplay with initiator methionine removal by methionine aminopeptidases and supported by extra translational evidence provided by Ribo-seq (included in the custom search).

**Figure 2:**
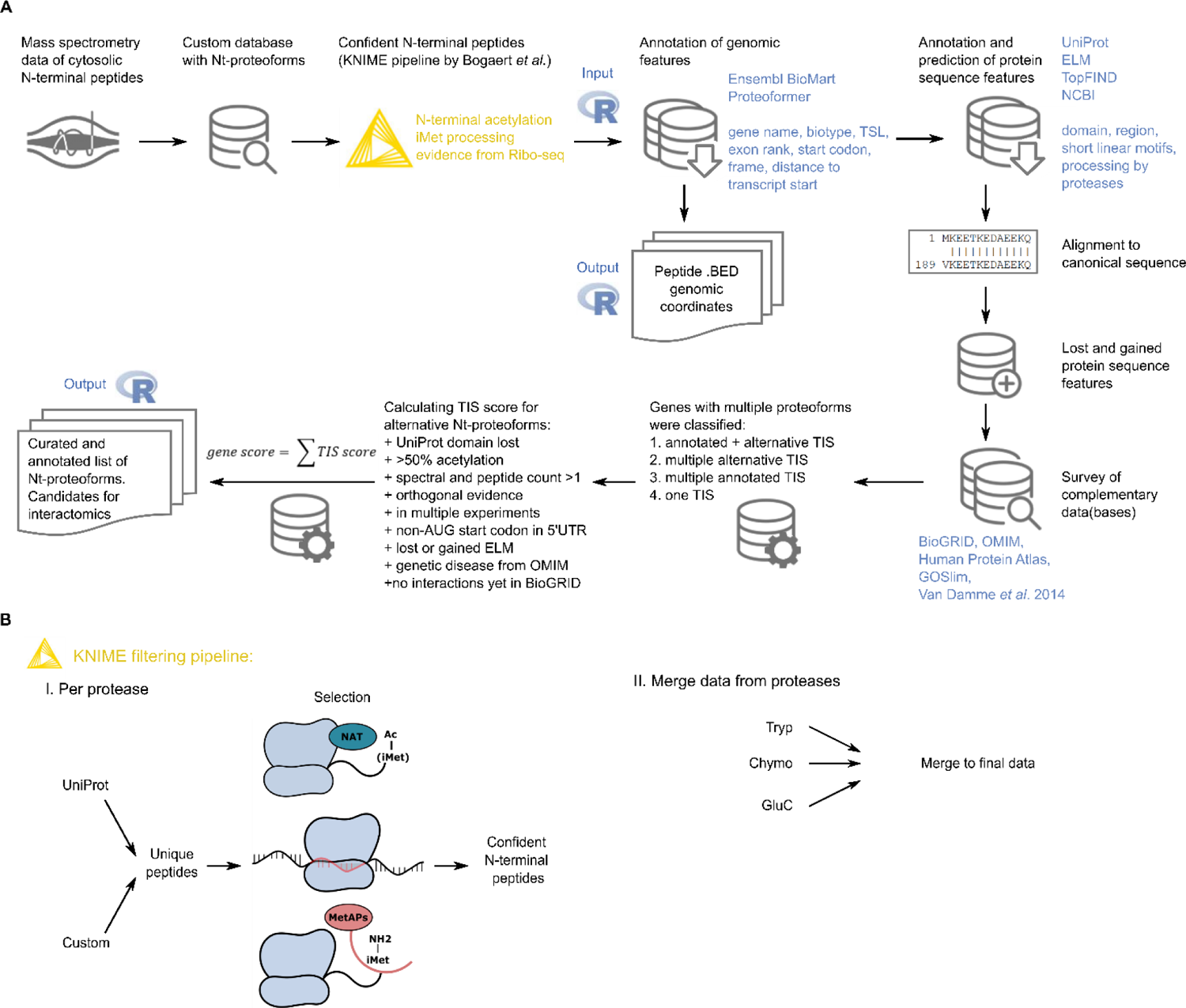
Selection of candidate Nt-proteoforms. A) The N-terminal proteomics data curation and analysis workflow included a KNIME pipeline for stringent filtering of Nt-peptides (see Bogaert et al. [43] and *Figure 2B*), followed by annotation of genomic and protein sequence features of candidate Nt-proteoforms using a custom R workflow. Based on the collected information, a TIS score and its derivative gene score were calculated to prioritize genes with alternative Nt-proteoforms for interactome validation. B) Overview of the Nt-peptide filtering workflow. Per protease, the data are merged and filtered to obtain unique peptide sequences before selection of confident N-terminal peptides originating from translation. This selection is done based on co-translational acetylation of a protein’s N-terminus, extra translational evidence by Ribo-seq and the presence or potential processing of the initiator methionine (iMet) by methionine aminopeptidases (MetAPs). Confident N-terminal peptides found for each protease-treated sample were merged to obtain a dataset of N-terminal peptides.

By this filtering strategy, we only retained confident N-terminal peptides pointing to database-annotated protein starts or N-terminal proteoforms, and assigned a confidence level to these peptides. As a last step, Nt-peptides identified by different proteases were merged into a final dataset (see **Figure 2**) of 3,306 unique Nt-peptides (see **Table 1** and **Supplementary Table S2**). We show that the combination of data from three different proteases increases the proteome coverage (see **Supplementary Figure S1.D**).

**Table 1:**
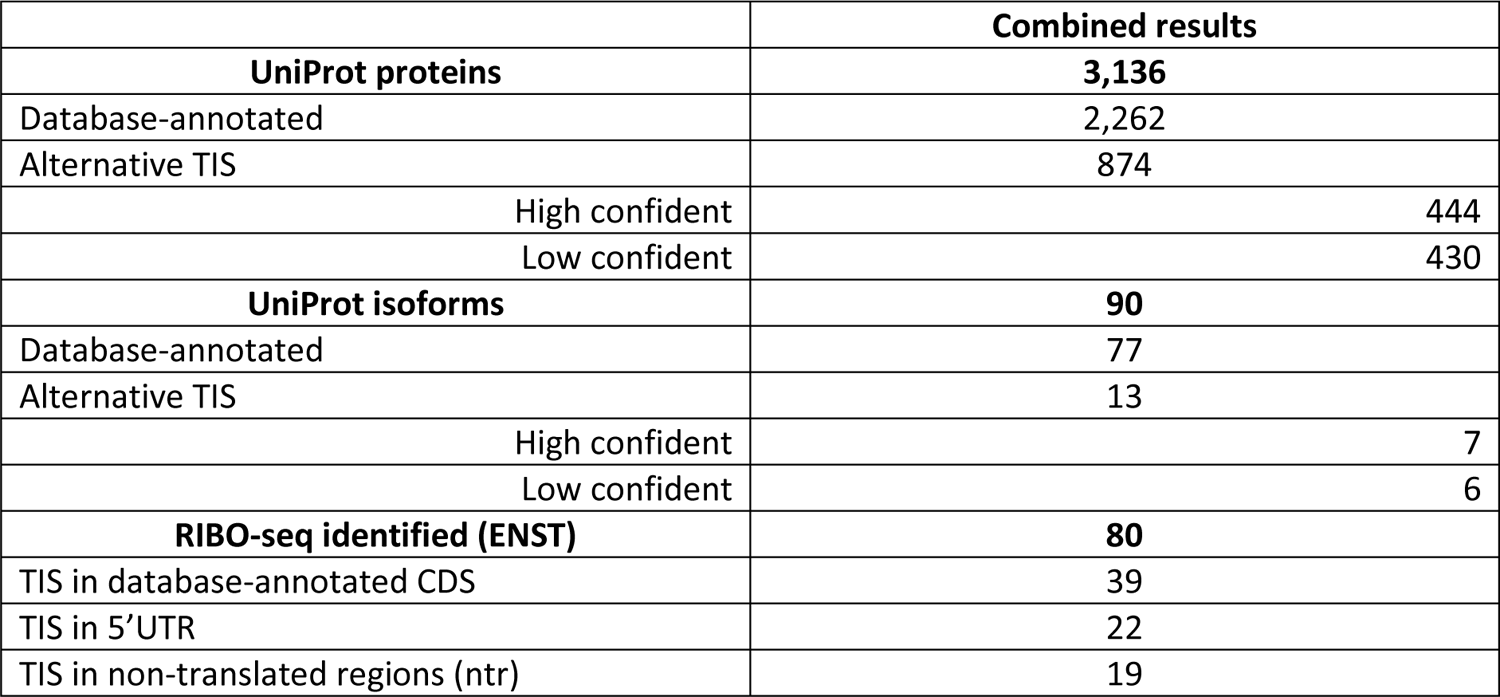
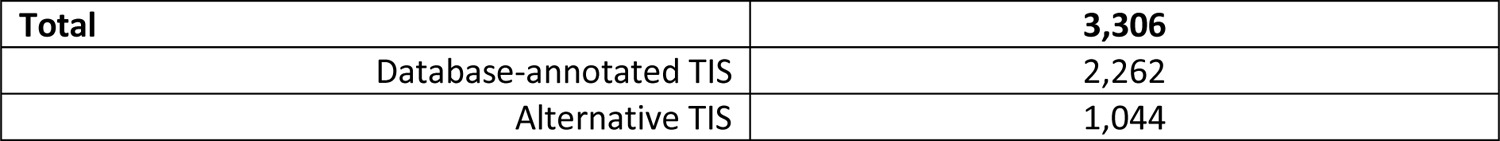
Overview of all identified N-terminal peptides after applying stringent selection. Nt-peptides are classified based on the type of protein sequence they are linked to, being a regular UniProt protein, a UniProt isoform or a RIBO-seq derived protein sequence. A distinction is made between database-annotated Nt-peptides (starting at position 1 or 2) and peptides with a start position beyond two, thus pointing to N-terminal proteoforms (alternative translation initiation start site (TIS)). Based on translational evidence, a confidence level is assigned (either low or high confident) to the identified Nt-proteoforms.

As can be seen in **Table 1** and **Supplementary Table S2**, the majority of identified peptides are known Nt-peptides listed in UniProt (2,262 or 68.4%), while the remaining 1,044 (31.6%) Nt-peptides point to potential (Nt-)proteoforms. From these potential Nt-proteoforms, 874 N-terminal peptides were identified that stem from translation starting at an internal site of a UniProt canonical protein. We also identified 90 Nt-peptides only matching a UniProt isoform and 80 Nt-peptides only matching an Ensembl entry (a proteoform translated from Ensembl transcript and derived from RIBO-seq evidence). Nt-peptides matching non-canonical accessions often originate from N-terminal proteoforms as these Ensembl and UniProt isoforms accessions include splice variants (with differences at the N-terminus) or the products of alternative translation initiation.

### 2. Selection of N-terminal proteoforms for interaction mapping

To trim down the catalogue of 1,044 N-terminal proteoforms to a more manageable set of pairs of N-terminal proteoforms and corresponding canonical proteins to test by Virotrap, we gathered extra information on the identified proteins to select confident and potentially interesting pairs.

As Virotrap is currently restricted to cytosolic proteins, it is important to verify the cytosolic localization of both the canonical protein and its N-terminal proteoform. Therefore, we generated a map of cytosolic proteins in HEK293T cells. Cytosolic extracts were prepared with 0.02% digitonin of HEK293T cells in triplicate and, following trypsin digestion and peptide pre-fractionation, LC-MS/MS analysis led to the identification of 3,045 proteins (**Supplementary Table S3**). Besides the GO terms added in the Perseus analysis, all identified proteins were submitted to the Retrieve/ID mapping tool on the UniProt website to evaluate their associated Gene Ontology Cellular Component (GOCC) terms and around 60% of these proteins contain the GO term “cytosol” (GA:0005829). When considering LFQ intensities of these cytosolic proteins, we see that they account for 86% of the sample. Note that, this map will be used for the selection of proteoform pairs.

To gather more gene and protein centered information, an R workflow was developed (see **Figure 2A**) to extract genomic information about the proteoforms (such as genomic coordinates, exons, start codon, frame, gene, biotype, TSL (transcript support levels)) and find protein sequence features (such as known domains and eukaryotic linear motifs, ELMs) that can be lost/gained by proteoforms of the same gene. We supplemented this information with disease association from the OMIM database, known interaction partners in BioGRID, whether or not the Nt-peptide was previously reported [5], and cytosolic localization reported in our own cytosolic map (see above), the Human Protein Atlas and gene ontology GOSlim annotation. Based on this extra information, including lost protein domains, different predicted linear motifs, the use of non-AUG start codons, presence of co-translational modifications, higher spectral and peptide counts, orthogonal evidence from UniProt, Ensembl or Ribo-seq, we built a scoring system considering the most relevant parameters listed for each peptide, called TIS scoring. TIS scores of the different Nt-peptides identified from the same gene were then combined into a gene score (see **Figure 2A**). Most importantly, when there is evidence that the detected peptide is not an N-terminal peptide resulting from translation but rather from processing (potential dipeptidase, signal/transit/pro-peptide processing or potential cleavage site of a protease), TIS scores drop to zero. The resulting list of Nt-peptides including all extra information and TIS/gene scores can be found in **Supplementary Table S4.** In total, 372 genes (corresponding to 868 Nt-peptides) have a gene score > 0 (see **Supplementary Table S4**, second tab).

In order to select pairs of canonical proteins and N-terminal proteoforms for interactome profiling, we developed a prioritization strategy based on protein localization, expression and other functional and structural parameters (see below). To gather more information about their expression, we examined if the identified N-terminal proteoforms are also expressed in other cell lines or tissues as this expands their biological significance. This analysis was done by exploiting Western blot data present in The Human Protein Atlas (HPA) for these 372 genes. To detect N-terminal proteoforms on WB, clearly the difference in molecular weights between the canonical protein and N-terminal proteoform needs to be sufficiently large to be detectable, and the epitope targeted by the antibody needs to be preserved in the proteoform, which reduced the list to 138 genes. For each of these, information on all tested antibodies (also unpublished ones) was extracted and antibodies with a WB score of two, which indicates the detection of a protein band of the predicted size (± 20%) with additional bands present, were selected. In total, data on 621 antibodies detecting the protein products of 136 genes were retrieved. Note that for two genes, no antibody information was present. For 143 of the 621 antibodies, extra bands were reported and the blots with these antibodies (detecting the protein products of 95 genes) were evaluated in more detail (online at www.proteinatlas.org, under antibodies and validation). For the protein products of 34 genes, an extra band corresponding to the size of the N-terminal proteoform(s) was detected, indicating a plausible expression of these Nt-proteoforms in other cell lines and/or tissues (see **Supplementary Table S5**). For example, a band corresponding to the Nt-proteoform of FNTA (40.4 kDa instead of 44.4 kDa, antibody CAB010149) was found in RT4 and U-251 mg cell lines, as well as in liver and tonsil tissue.

To further evaluate the expression of Nt-proteoforms in healthy human tissues, we re-analyzed public proteomics data of the draft map of the human proteome developed by the Pandey group [55]. The use of ionbot [56] and a custom-build protein sequence database (composed of UniProt and Ribo-seq derived proteoforms) led to 9,151,086 peptide to spectrum matches (PSMs). Further filtering and aggregation of the data was performed in R, leading to 8,501,009 filtered PSMs and 2,789,079 unique peptides belonging to 26,159 proteoforms. Of the 3,306 proteoforms identified by N-terminal COFRADIC, 897 were found expressed in human tissues and supported by an N-terminal peptide identified at a matching start position. Among these 897 proteins, 24 are non-canonical Nt-proteoforms, thus proteoforms with peptides that do not match a canonical UniProt protein (**Figure 3** panel A, **Supplementary Table S6**). Differential expression analysis across tissues using normalized spectral abundance factors (NSAF) indicated that 582 out of 897 proteoforms matching N-terminal COFRADIC data had a significant tissue-dependent expression profile, including three non-canonical proteoforms of two genes, namely TPM3 and EPB41L3 (**Figure 3** panel B and C). These data confirm that Nt-proteoforms are not only expressed in histologically healthy human tissues, but also display tissue specificity (**Figure 3A**) or different tissue expression profile when comparing Nt-proteoforms of the same gene (**Figure 3B-C**).

**Figure 3:**
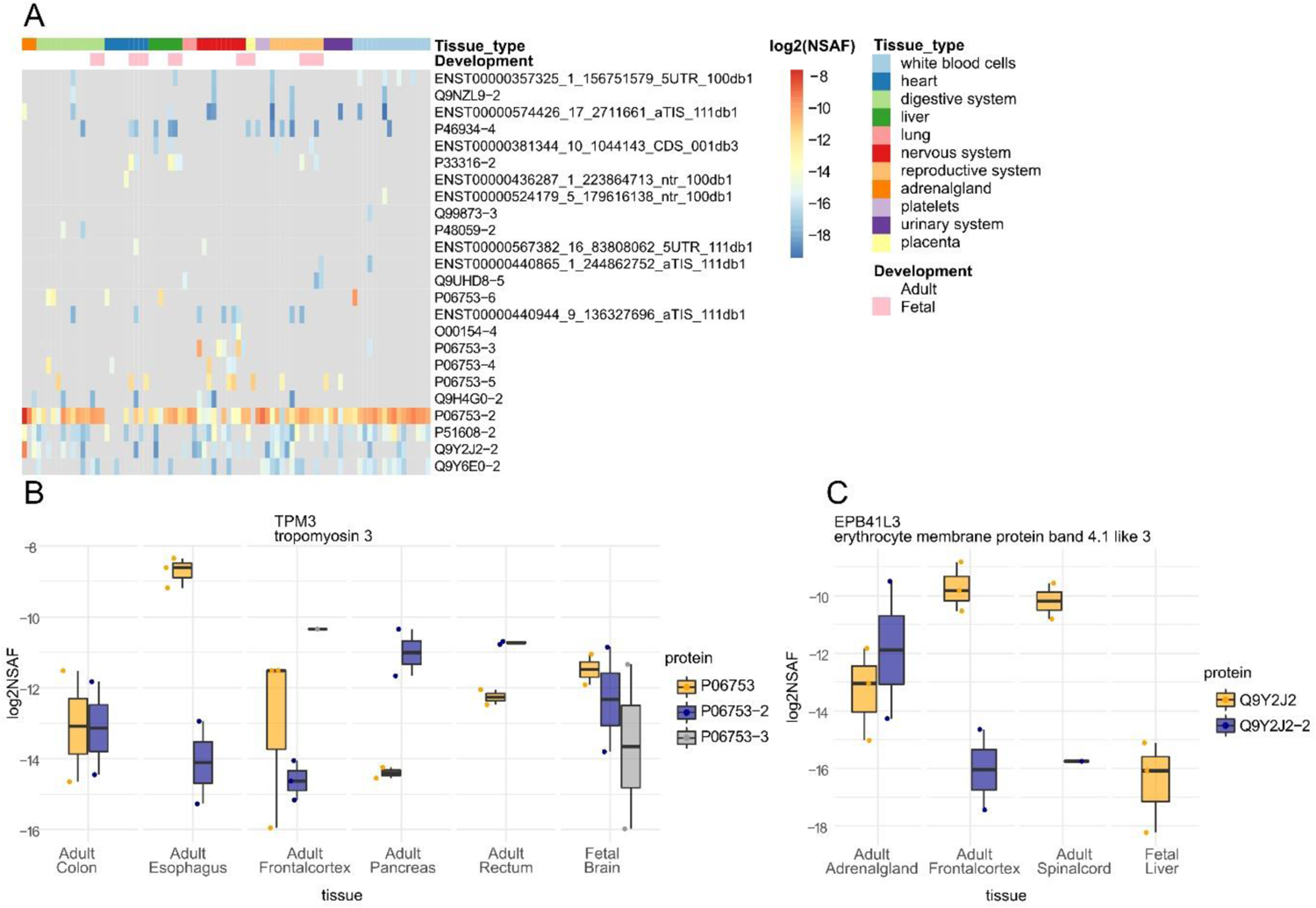
Tissue expression of Nt-proteoforms. A. Heatmap presenting log2NASF expression values of 24 non-canonical proteoforms across healthy human tissues from the Pandey dataset [55]. Nt-proteoforms from the same gene (TPM3 in panel B and EPB41L3 in panel C) show distinct profile of tissue expression. All presented proteoforms were found significant in our differential expression analysis (adjusted p-value ≤ 0.05).

For the final selection of candidates for interactome profiling by Virotrap we applied additional criteria for prioritization. First, we prioritized proteoforms with a loss of a known domain or eukaryotic linear motif (ELM). Then, we prioritized proteoforms with a considerable length of truncation or extension (>20 amino acids or >50 amino acids for proteins over 700 amino acids). Next, we prioritized proteoforms suited for Virotrap, being cytosolic proteoforms (by checking our cytosolic map and the subcellular localizations listed in UniProt and HPA), and non-structural proteins and enzymes over structural proteins. Finally, we prioritized for known disease-associated proteins and proteoforms over novel proteins (e.g. proteins from out-of-frame translation) and considered the results of the HPA-WB analysis.

As a final result, we report 85 proteoform pairs meeting several selection criteria, of which the 22 highest ranking (best scoring over all the criteria) were selected for analysis by Virotrap (see **Table 2** and **Supplementary Table S4** (third worksheet)). Note that not for all proteins the canonical N-terminus was identified. This was the case for CAST, CSDE1, SPAST and UBXN6. In these cases, verification of the cytosolic localization is important as it is known that N-terminal proteoforms and their canonical proteins can have different subcellular localizations [14]. Note also that the results of the protein originating from a presumed non-translated region (NTR, ACTB pseudogene 8) have been published before [45] and will thus not be further discussed here. When checking the 175 Nt-proteoforms corresponding to 85 genes considered for Virotrap experiments in the re-analyzed public proteomics data of the draft map of the human proteome, we found 100 Nt-proteoforms expressed in human tissues (see **Supplementary Table S6** column “virotrap_intersect”).

**Table 2:** List of the proteins that were selected for interactome analysis. For each protein, information about the canonical protein is provided (UniProt protein accession and gene name) and about the N-terminal proteoform(s) concerning the length of the truncation or extension. In the last column, differences between the canonical protein and the N-terminal proteoforms with regards to domains, motifs or ELMs are listed.

| Protein | Gene | Proteoform | Lost or gained domains and motifs |
| --- | --- | --- | --- |
| Q12904<br>Aminoacyl tRNA synthase complex-interacting multifunctional protein 1<br>Length: 312 AA | AIMP1 | Known UniProt isoform Q12904-2; N-terminal extension of 24 AA | Gain of three linear motifs according to ELM: binding motif for UBA3 adenylation (AA 17-24), SH3 ligand (AA 4-10) and SPAK-OSR1 docking motif (AA 14-18) |
| Q9HB71<br>Calcyclin-binding protein<br>Length: 228 AA | CACYBP | Known UniProt isoform Q9HB71-3; N-terminal truncation of 43 AA. | Partial loss of region for interaction with SIAH1 (AA 2-80) |
| Q14444<br>Caprin-1<br>Length: 709 AA | CAPRIN1 | High confident N-terminal proteoform; N-terminal truncation of 116 AA | Loss of coiled coil domain (60-94) |
| P20810<br>Calpastatin<br>Length: 708 | CAST | The identified N-terminal peptide cannot differentiate which proteoform was identified: known UniProt isoforms P20810-4 (missing AA 9-30 and AA 44-62) or P20810-8 only missing AA 9-30. | - |
| Q9Y281<br>Cofilin-2<br>Length: 166 AA | CFL2 | Known UniProt isoform Q9Y281-3; N-terminal truncation of 17 AA | Partial loss of the region for interaction with CSRP3 (AA 2-55) and partial loss of ADF-H domain (AA 4-153). |
| O75535<br>Cold shock domain-containing protein E1<br>Length: 798 AA | CSDE1 | High confident N-terminal proteoform; N-terminal truncation of 176 AA | Loss of CDS1 domain (AA 26-87) and loss of majority of the CSD2 domain (AA 136-179) |
| P60842<br>Eukaryotic initiation factor 4A-I<br>Length: 406 | EIF4A1 | High confident N-terminal proteoform; N-terminal truncation of 211 AA | Partial loss of helicase ATP-binding domain (63-234), loss of Q motif (AA 32-60) and loss of DEAD box motif (AA 182-185) |
| P49354<br>Protein farnesyltransferase /geranylgeranyltransferase type-1 subunit alpha<br>Length: 379 AA | FNTA | High confident N-terminal proteoform; N-terminal truncation of 38 AA | Loss of a Pro-rich region (AA 22-31) |
| Q7Z434<br>Mitochondrial antiviral-signaling protein<br>Length: 540 AA | MAVS | Known UniProt isoform Q7Z434-4; N-terminal truncation of 141 AA | Loss of CARD domain (AA 10-77), loss of region for interaction with NLRX1 (AA 10-77), partial loss of Pro-rich region (AA 103-153) and region for TRAF2 interaction becomes outer N-terminal |
| Q9H074<br>Polyadenylate-binding protein-interacting protein 1<br>Length: 479 AA | PAIP1 | Known UniProt isoform Q9H074-3; N-terminal truncation of 112 AA | Loss of Gly-rich region (AA 10-36), loss of Pro-rich region (AA 45-98) and PABPC interaction motif 2 becomes N-terminal (AA 116-143) |
| Q14558<br>Phosphoribosyl pyrophosphate synthase-associated protein 1<br>Length: 356 AA | PRPSAP1 | Known UniProt isoform Q14558-2; N-terminal extension of 29 AA | Gain of 11 linear motifs according to ELM |
| Q86TP1<br>Exopolyphosphatase PRUNE<br>Length: 453 AA | PRUNE | The identified N-terminal peptide cannot differentiate which proteoform was identified: known UniProt isoforms Q86TP1-3 (missing AA 1-182) and Q86TP1-5 (missing AA 1-182 and AA 259-311) | Loss of DHH motif (AA 106-108) |
| P49023<br>Paxillin<br>Length: 591 | PXN | Known UniProt isoform P49023-4; N-terminal truncation of 133 AA and also missing AA 278-311 | Loss of LD motif (AA 3-15) and loss of Pro-rich region (AA 46-53) |
| P54136<br>Arginine-tRNA ligase,<br>cytoplasmic<br>Length : 660 AA | RARS | Known UniProt Isoform P54136-2; N-terminal truncation of 72 AA | Loss of a region that could be involved in the assembly of the multisynthetase complex (AA 1-72) |
| Q9UBP0<br>Spastin<br>Length: 616 AA | SPAST | The identified N-terminal peptide cannot differentiate which proteoform was identified: known UniProt isoforms Q9UBP0-3 (missing AA 1-86) or Q9UBP0-4 (missing AA 1-86 and AA 197-228) | Loss of region for interaction with ATL1 (AA 1-80), loss of region needed for nuclear localization (AA 1-50), loss of region for interaction with SSNA1 and microtubules (AA 50-87), loss of NLS (AA 4-11), loss on nuclear export signal (AA 59-67), partial loss of region for interaction with RTN1 (AA 1-300) and partial loss of region for midbody localization (AA 1-194) |
| Q99576<br>TSC22 domain family protein 3<br>Length: 134 AA | TSC22D3 | High confident N-terminal proteoform; N-terminal truncation of 57 AA. | Almost complete loss of AP1-binding region |
| Q9BSL1<br>Ubiquitin-associated domain-containing protein 1<br>Length: 405 AA | UBAC1 | High confident N-terminal proteoform; N-terminal truncation of 102 AA | Loss of ubiquitin-like domain (AA 14-98) |
| P61081<br>NEDD8-conjugating enzyme Ubc12<br>Length: 183 AA | UBE2M | High confident N-terminal proteoform; N-terminal extension of 42 AA. | Gain of a Caspase cleavage motif (AA 14-18), gain of a Glycosaminoglycan attachment site (AA 13-16 and 35-38), gain of IAP-binding motif (AA 1-4 and 17-21) and gain of a TRAF2 binding site (AA 3-6) |
| Q9BZV1<br>UBX domain containing protein 6<br>Length: 441 AA | UBXN6 | Known UniProt isoform Q9BZV1-2; N-terminal truncation of 53 AA | Loss of region for interaction with LMAN1 (AA 1-10) and VCP/p97-interacting motif (VIM) becomes N-terminal, with 2 AA removed (AA 51-63) |
| P09936<br>Ubiquitin carboxyl-terminal hydrolase isozyme L1<br>Length: 223 AA | UCHL1 | Two high confident N-terminal proteoforms: N-terminal truncations of 5 or 11 AA | Proteoform 1: ubiquitin interaction domain (AA 5-11) becomes outer N-terminal<br>Proteoform 2: loss of ubiquitin interaction domain (AA 5-11) |
| Q8TCF1<br>AN1-type zinc finger protein 1<br>Length: 268 | ZFAND1 | Known UniProt isoform Q8TCF1-2; N-terminal truncation of 107 AA | Loss of AN1-type 1 Zinc finger domain (7-58) and loss of the majority of AN1-type 2 Zinc finger domain (AA 61-110) |
| ENST00000403258<br>ACTB pseudogene 8<br>Length: 146 AA | NTR protein | High confident N-terminal proteoform; N-terminal truncation of 11 AA | ELM predicts several motifs and domains for this protein. |

### 3. Interaction profiling of Nt-proteoforms and their canonical counterparts

Mapping of the protein complex a protein is engaged in is a frequently used approach to gain insights into the processes and pathways that protein is involved in (the “guilt-by-association” approach) [35]. However, most interactome studies are restricted to studying the canonical protein, but as (N-terminal) proteoforms can have altered functions, including these proteoforms in interactomics studies would help us to better understand their function and how this function is possibly different from that of the canonical protein. For interactome analysis, Virotrap is used. In short, a bait protein is fused to the C-terminus of GAG, leading to the recruitment and multimerization of the GAG-bait fusion protein at the plasma membrane followed by subsequent budding of VLPs from the cells. As the bait is coupled to GAG this allows for co-purification of bait-associated protein partners by trapping them into VLPs. Purification of the VLPs relies on co-expressing FLAG-tagged and untagged VSV-G, presented as trimers on the surface of VLPs, allowing for efficient antibody-based purification of the VLPs

To study the interactomes of pairs of canonical proteins and N-terminal proteoforms, we designed the following strategy (**Figure 4**). First, both canonical proteins (full length, further referred to as FL) and N-terminal proteoforms (referred to as PR) were cloned into pMET7-GAG-bait plasmids where they are N-terminally fused to the GAG protein. This led to 45 pMET7-GAG-bait constructs of 21 genes with for each gene one FL bait and one or two PR baits (**Supplementary Figure S2.A**). In an initial Virotrap screen, bait expression was first tested on Western blots (**Figure 4.A**) to evaluate if baits were well expressed, FL and PR baits had comparable expression levels and if they were efficiently recruited into VLPs. Note that similar expression levels of the control bait (eDHFR), FL and PR are desired for statistical analysis of the LC-MS/MS data (**Figure 4.A**). Baits that were not recruited into VLPs were left out for further analysis, which was the case for CAST and RARS. For CAST, both FL and PR were not pulled into VLPs, while for RARS only the FL was pulled into the VLPs (**Figure 4.B,** uncropped WB are shown in **Supplementary Figure S2.B**), not allowing for comparative analysis. All other baits (e.g. PAIP1, **Figure 4.B**) were clearly detected in the cell lysates and in the VLPs.

**Figure 4:**
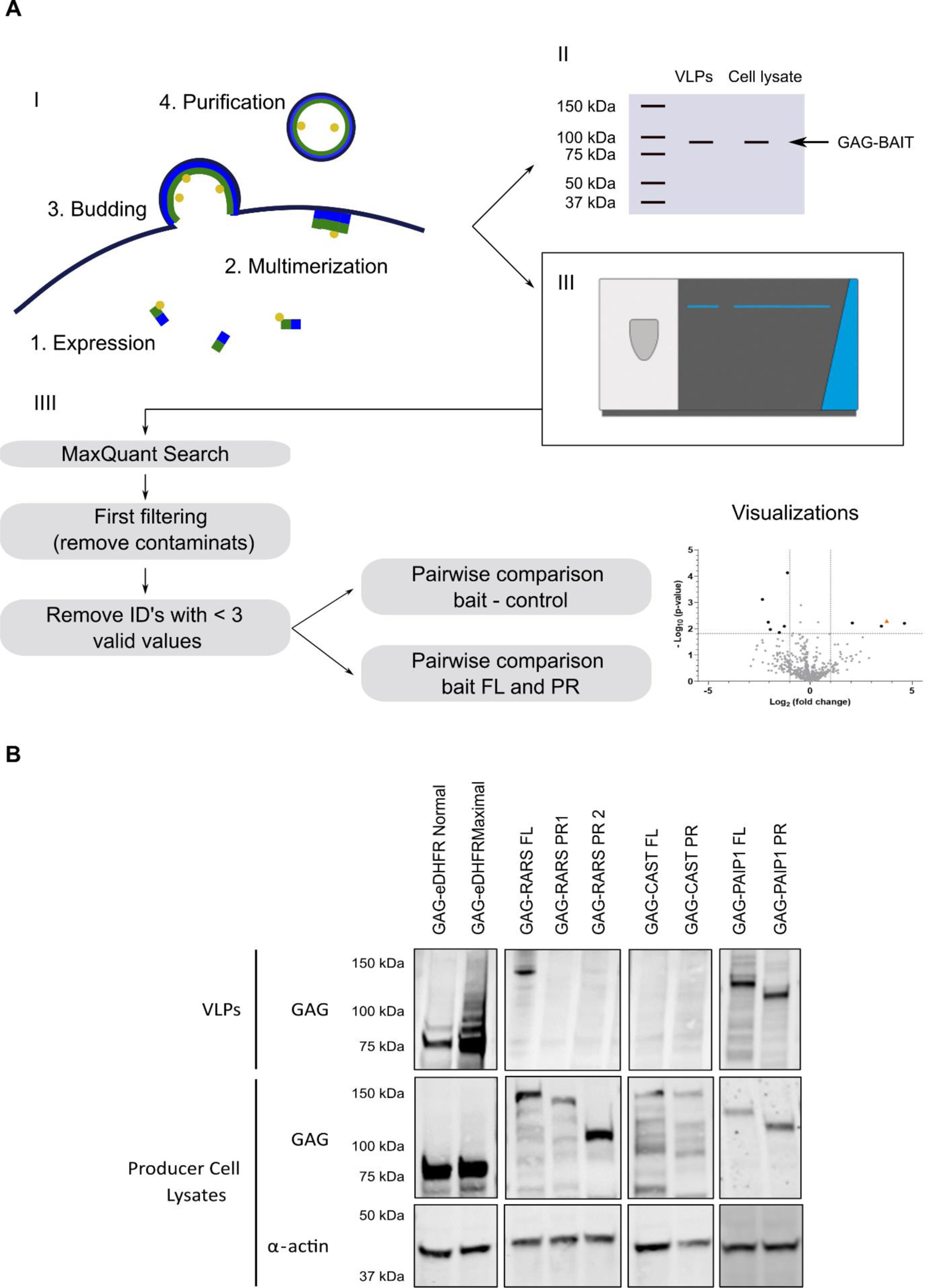
Overview of the Virotrap workflow. A) Protein complex purification by Virotrap and data analysis approach I. GAG-bait fusion proteins were expressed in HEK293T cells and, following multimerization at the plasma membrane, GAG-bait fusion proteins and their interacting proteins are trapped in viral-like particles (VLPs) that bud from the cells. These VLPs are subsequently purified from the media and prepared for either WB or LC-MS/MS analysis. II. As an initial screen, all baits are analyzed by WB to validate proper expression and recruitment into VLPs. III. Baits that are successfully recruited into VLPs (as determined by WB) are suited for further experiments. For LC-MS/MS, three biological replicates of all baits were performed. Baits were divided into manageable sets of one control bait and maximally three pairs of FL and PR. III. The LC-MS/MS data were searched with MaxQuant’s Andromeda search engine against the human proteome (supplemented with the sequences of the proteins expressed for Virotrap such as VSV-G and with only the shortest sequence of the baits), filtered (to remove reversed matches, proteins only identified by site and contaminants) and the LFQ intensities were transformed. Afterwards, samples were grouped and identified proteins were filtered on three valid values in at least one group and subsequently imputed. Potential interaction partners were identified by pairwise comparison with control samples and functional differences between FL and PR were highlighted by pairwise comparisons between FL and PR. Data can be visualized in different ways, e.g. a volcano plot. B) Western blot results of initial Virotrap screens for the detection of expression and recruitment into the VLPs of GAG-bait fusion proteins. HEK293T cells were transfected with GAG-bait constructs. Additional co-transfection of VSV-G/FLAG-VSV-G expression constructs allowed VLP purification, which was followed by direct on-bead lysis and analysis by Western blotting using anti-GAG (bait expression levels and particles) and anti-α-actin antibodies (as loading control for cell lysates). Results of VLPs and producer cell lysates are shown. For GAG-eDHFR and GAG-PAIP1 (both FL and PR), a clear band can be detected at the desired molecular weight in both VLPs and cells. For RARS, all constructs seem well expressed in the cells, but only RARS FL is recruited into VLPs. CAST is found only weakly expressed in cells and is not pulled into VLPs. Uncropped gel images and molecular weight markers are shown in Supplementary *Figure S2*.B.

For LC-MS/MS analysis, we divided the baits in manageable sets including control samples and maximally three pairs of FL and PR. Here, baits with similar expression levels were combined and triplicate experiments for all baits were performed. In total, baits were divided into seven sets (see Materials and Methods section). To obtain specific interaction partners of the FL or PR from the lists of identified proteins, their interactomes were compared with that of the control baits. To evaluate possible functional differences between FL and PR, the identified proteins using the FL baits were directly compared with those from the PR baits (**Figure 4.B**).

Triplicate Virotrap experiments were performed for the protein products of 20 genes (42 proteoforms) taking along GAG-eDHFR as control. In the following, we illustrate the employed data analysis strategy for MAVS, and this strategy is applied on all baits in the same way. MAVS is annotated as a mitochondrial antiviral-signaling protein for which a N-terminal proteoform starting at methionine-142 was identified, which is a known UniProt isoform. This proteoform loses a CARD domain, which is required for interaction with NLRX1, and thereby the region required for interaction with TRAF2 becomes outer N-terminal (**Figure 5.A**).

**Figure 5.**
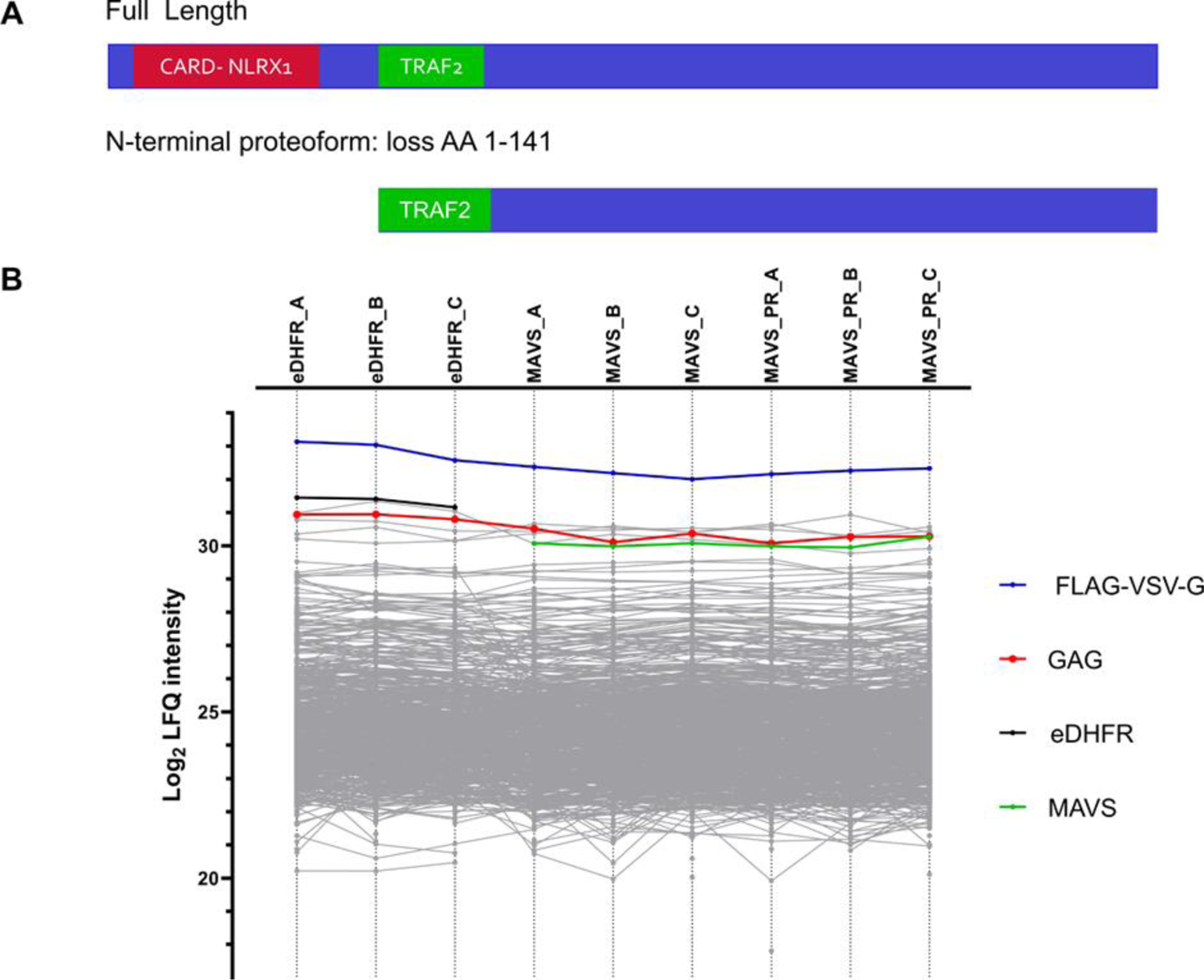
Comparisons of bait intensities. A) Schematic representation of the differences in domains for full-length MAVS and its N-terminal proteoform. B) Profile plot showing the log_2_ transformed label free quantification (LFQ) intensities for the different identified proteins in the MAVS Virotrap experiment. LFQ intensities for every replicate are shown before imputation of missing values for statistical analysis.

MaxQuant searches of the experimental set including MAVS FL and PR, led to the identification of 2,134 unique proteins. Further data analysis was handled in Perseus, and after removal of contaminants, reversed proteins and proteins only identified by site, 1,997 proteins remained. The protein LFQ intensities were log_2_ transformed and replicates were grouped. Proteins identified in less than three samples in at least one group were removed, leading to a final set of 842 proteins. The missing values were imputed using imputeLCMD. As already mentioned, the bait levels are ideally very similar allowing a straightforward comparison of the levels of interaction partners or commonly co-purified proteins between the different bait interactomes [65]. Therefore, as a first check, bait intensities were visualized in a profile plot (before imputation) and both MAVS FL and PR had comparable intensities however, they both are less intense compared to the control (eDHFR) intensities (**Figure 5.B**).

For 13 baits, CACYBP, CAPRIN1, CSDE1, EIF4A1, FNTA, MAVS, PAIP1, PRPSAP1, UBAC1, UBE2M, UBXN6, UCHL1 and ZFAND1, FL and PR(s) showed similar expression levels. For the seven other baits, AIMP1, CFL2, PRUNE, PXN, NTR, SPAST and TSC22D3, a difference in expression levels of at least two-fold was found between the FL and PR, hampering conclusive statistical analysis (**Supplementary Figure S3**). Data of all baits are here reported. Subsequently, all baits were pairwise tested against the eDHFR control samples to identify their candidate interaction partners at an FDR ≤ 0.01, which resulted in 43 candidate interaction partners for MAVS FL and 48 for MAVS PR (**Table 3**). We compared such lists of potential interaction partners with known interaction partners listed in BioGRID [50], STRING [51] and IntAct [52]. Amongst the candidate MAVS interaction partners, we found BAG6, IFIT1, IFNB1 and TRAF2 (only reported for the FL), which are known interaction partners of MAVS. Of the 43 candidate interaction partners of the canonical protein, 39 (90.7%) are also found using the N-terminal proteoform as bait. Four proteins seem to be potential interaction partners for full-length MAVS, ACLY, EEF2, PKM and TRAF2, while nine proteins, ACTL6A, CAD, DPYSL2, PGM1, PKRKACB, PLK1, PRKDC, PYGL and SEPT7, seem to be specific for the shorter MAVS proteoform, possibly pointing to functional diversities of these two MAVS variants.

**Table 3:**
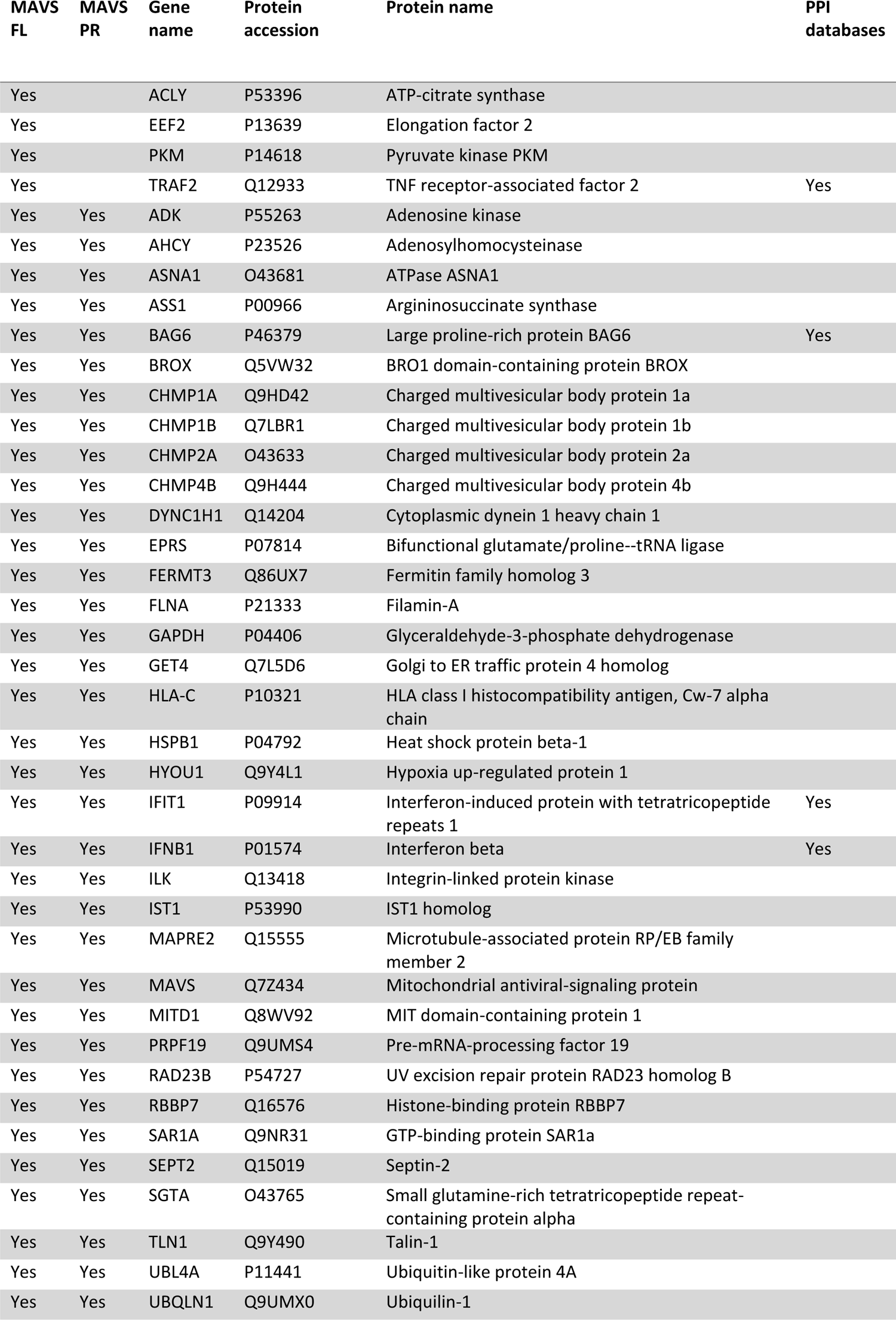
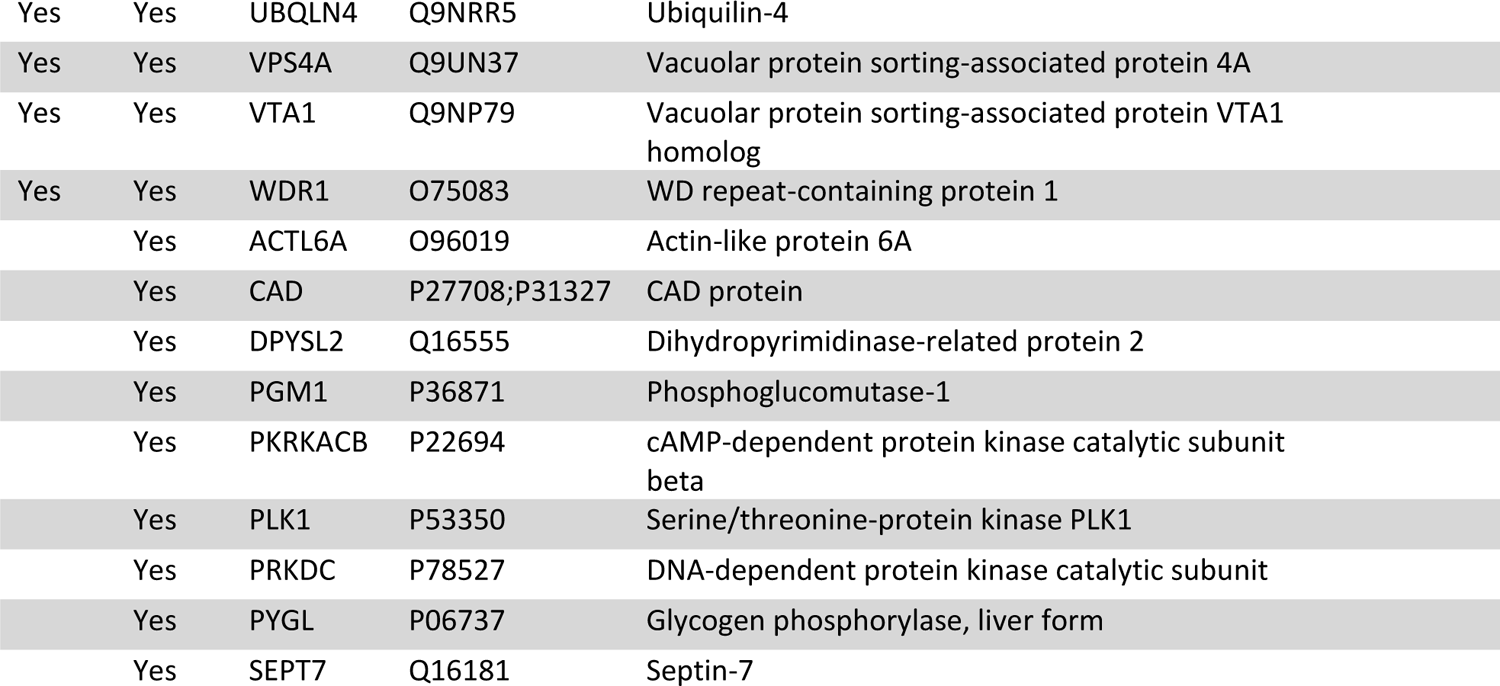
List of MAVS candidate interaction partners. MAVS FL and PR Virotrap experiments were performed in triplicate and data were compared with eDHFR control samples. The first two columns indicate whether a protein was identified as a candidate MAVS FL or PR interaction partner, the last column indicates whether a protein is a known MAVS interaction partner as listed in the BioGRID, STRING and/or IntAct databases.

| MAVS FL | MAVS PR | Gene name | Protein accession | Protein name | PPI databases |
| --- | --- | --- | --- | --- | --- |
| Yes |  | ACLY | P53396 | ATP-citrate synthase |  |
| Yes |  | EEF2 | P13639 | Elongation factor 2 |  |
| Yes |  | PKM | P14618 | Pyruvate kinase PKM |  |
| Yes |  | TRAF2 | Q12933 | TNF receptor-associated factor 2 | Yes |
| Yes | Yes | ADK | P55263 | Adenosine kinase |  |
| Yes | Yes | AHCY | P23526 | Adenosylhomocysteinase |  |
| Yes | Yes | ASNA1 | O43681 | ATPase ASNA1 |  |
| Yes | Yes | ASS1 | P00966 | Argininosuccinate synthase |  |
| Yes | Yes | BAG6 | P46379 | Large proline-rich protein BAG6 | Yes |
| Yes | Yes | BROX | Q5VW32 | BRO1 domain-containing protein BROX |  |
| Yes | Yes | CHMP1A | Q9HD42 | Charged multivesicular body protein 1a |  |
| Yes | Yes | CHMP1B | Q7LBR1 | Charged multivesicular body protein 1b |  |
| Yes | Yes | CHMP2A | O43633 | Charged multivesicular body protein 2a |  |
| Yes | Yes | CHMP4B | Q9H444 | Charged multivesicular body protein 4b |  |
| Yes | Yes | DYNC1H1 | Q14204 | Cytoplasmic dynein 1 heavy chain 1 |  |
| Yes | Yes | EPRS | P07814 | Bifunctional glutamate/proline--tRNA ligase |  |
| Yes | Yes | FERMT3 | Q86UX7 | Fermitin family homolog 3 |  |
| Yes | Yes | FLNA | P21333 | Filamin-A |  |
| Yes | Yes | GAPDH | P04406 | Glyceraldehyde-3-phosphate dehydrogenase |  |
| Yes | Yes | GET4 | Q7L5D6 | Golgi to ER traffic protein 4 homolog |  |
| Yes | Yes | HLA-C | P10321 | HLA class I histocompatibility antigen, Cw-7 alpha chain |  |
| Yes | Yes | HSPB1 | P04792 | Heat shock protein beta-1 |  |
| Yes | Yes | HYOU1 | Q9Y4L1 | Hypoxia up-regulated protein 1 |  |
| Yes | Yes | IFIT1 | P09914 | Interferon-induced protein with tetratricopeptide repeats 1 | Yes |
| Yes | Yes | IFNB1 | P01574 | Interferon beta | Yes |
| Yes | Yes | ILK | Q13418 | Integrin-linked protein kinase |  |
| Yes | Yes | IST1 | P53990 | IST1 homolog |  |
| Yes | Yes | MAPRE2 | Q15555 | Microtubule-associated protein RP/EB family member 2 |  |
| Yes | Yes | MAVS | Q7Z434 | Mitochondrial antiviral-signaling protein |  |
| Yes | Yes | MITD1 | Q8WV92 | MIT domain-containing protein 1 |  |
| Yes | Yes | PRPF19 | Q9UMS4 | Pre-mRNA-processing factor 19 |  |
| Yes | Yes | RAD23B | P54727 | UV excision repair protein RAD23 homolog B |  |
| Yes | Yes | RBBP7 | Q16576 | Histone-binding protein RBBP7 |  |
| Yes | Yes | SAR1A | Q9NR31 | GTP-binding protein SAR1a |  |
| Yes | Yes | SEPT2 | Q15019 | Septin-2 |  |
| Yes | Yes | SGTA | O43765 | Small glutamine-rich tetratricopeptide repeat-containing protein alpha |  |
| Yes | Yes | TLN1 | Q9Y490 | Talin-1 |  |
| Yes | Yes | UBL4A | P11441 | Ubiquitin-like protein 4A |  |
| Yes | Yes | UBQLN1 | Q9UMX0 | Ubiquilin-1 |  |

|  |  |  |  |  |
| --- | --- | --- | --- | --- |
| Yes | Yes | UBQLN4 | Q9NRR5 | Ubiquilin-4 |
| Yes | Yes | VPS4A | Q9UN37 | Vacuolar protein sorting-associated protein 4A |
| Yes | Yes | VTA1 | Q9NP79 | Vacuolar protein sorting-associated protein VTA1 homolog |
| Yes | Yes | WDR1 | O75083 | WD repeat-containing protein 1 |
|  | Yes | ACTL6A | O96019 | Actin-like protein 6A |
|  | Yes | CAD | P27708;P31327 | CAD protein |
|  | Yes | DPYSL2 | Q16555 | Dihydropyrimidinase-related protein 2 |
|  | Yes | PGM1 | P36871 | Phosphoglucomutase-1 |
|  | Yes | PKRKACB | P22694 | cAMP-dependent protein kinase catalytic subunit beta |
|  | Yes | PLK1 | P53350 | Serine/threonine-protein kinase PLK1 |
|  | Yes | PRKDC | P78527 | DNA-dependent protein kinase catalytic subunit |
|  | Yes | PYGL | P06737 | Glycogen phosphorylase, liver form |
|  | Yes | SEPT7 | Q16181 | Septin-7 |

For all baits tested in Virotrap, besides AIMP1, we were able to detect known interaction partners among the candidate interaction partners, validating our approach. All identified candidate interaction partners are listed in **Supplementary Table S7**. In general, we found high overlaps between the interactomes of paired proteoforms and, on average, 66.7 % of the candidate interaction partners of one proteoform were also found for the other proteoform, indicating that most interactors seem to be shared by the different proteoforms. For example, for cases such as UBXN6 (**Figure 6**top row left panel), the majority of candidate interaction partners is shared between the proteoforms, yet for each proteoform, unique interactors were found. On the other hand, for some other bait proteoform pairs, the overlap between interactors is quite limited, as shown for PRUNE (**Figure 6** top row middle panel) where only 25% of the candidate interaction partners are shared. On the contrary, for UBAC1 (**Figure 6**, top row right panel), while the N-terminal proteoform has lost several interactors, it but does not seem to engage in other protein-protein interactions.

**Figure 6:**
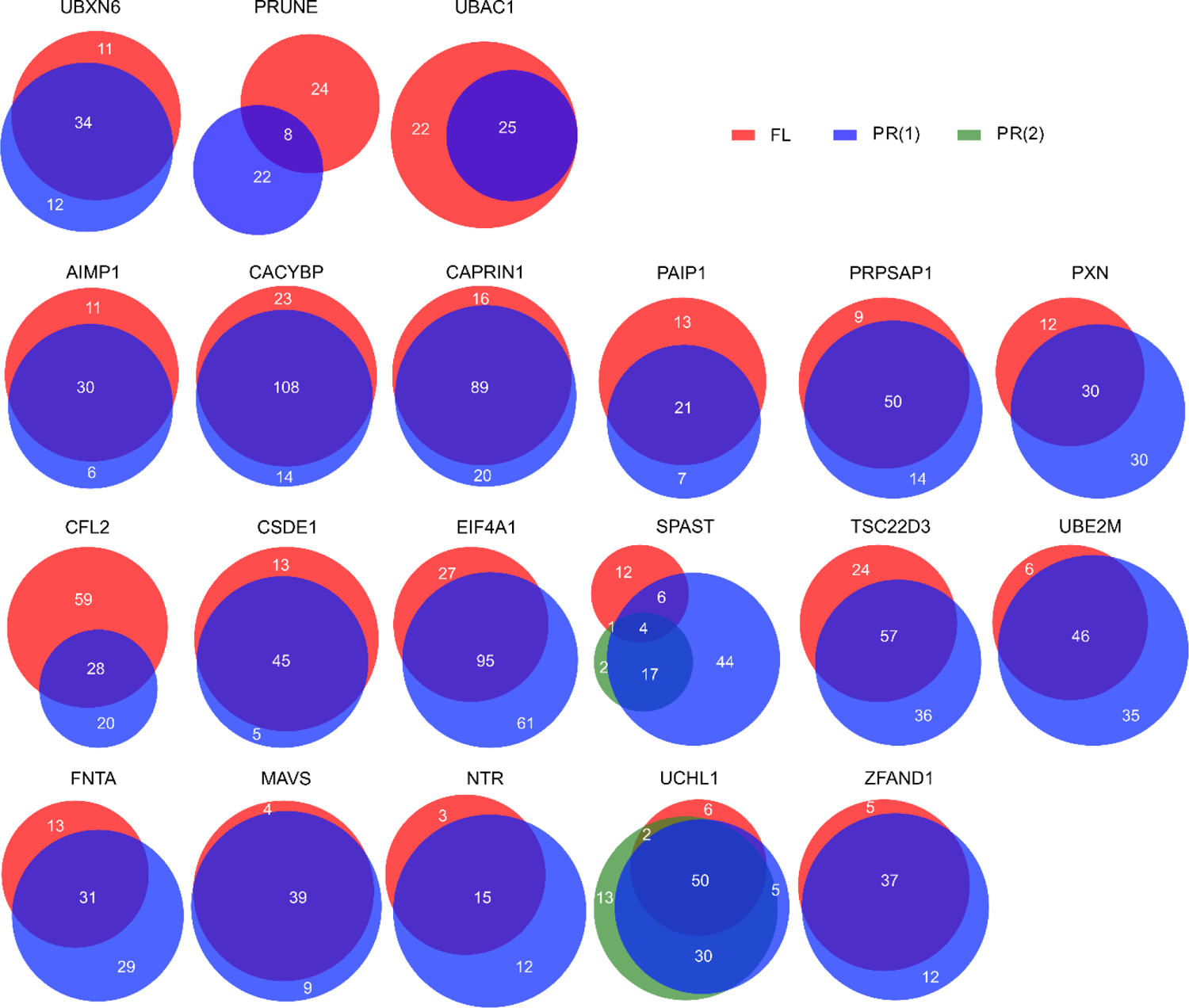
Overview of the overlap between the candidate interaction partners identified for the FL and the PR baits. The three baits on the top row show three different scenarios of differences between the interactome identified for the canonical protein and the N-terminal proteoform. The overlap between FL and PR of all other baits are shown.

As mentioned above, differences in candidate interaction partners between the canonical protein and N-terminal proteoform may point to functional diversities between protein variants. We aimed to study such differences in more detail by a direct pairwise comparison between the FL and PR interactomes, and withheld proteins with an at least two-fold change in levels (also correcting for differences in bait levels, see methods section) and with a pairwise test adjusted p-value (FDR) ≤ 0.05. Further, only proteins already listed as candidate interactors upon comparing with the interactomes of the control samples or already known as interaction partners as listed by in BioGRID [50], STRING [51] and IntAct [52] were retained.

The results of all pairwise comparisons between the different proteoforms of all baits are listed in **Supplementary Table S8** and visualized in **Supplementary Figure S3**. For all proteins we report differences between the FL and PR interactomes, and list if a prey was previously reported as a candidate interaction partner. In some comparisons, we report GAG amongst the significant proteins at an FDR ≤ 0.05. However, except for SPAST FL vs PR1 and SPAST PR1 vs PR2 interactomes, GAG is removed when also filtering on fold-change. We hypothesize that GAG might pop up due to differences in expression between FL and PR baits, but by our additional filtering on fold-change we removed GAG, showing the necessity of this additional filtering step to retain reliable differences and not differences only due to experimental variations. Along with GAG, for some cases, we report known GAG interaction partners amongst the significant proteins, but mostly not with a sufficiently large fold-difference, which illustrates these double filtering steps as a valid way for identifying proteoform-specific interactors.

### 4. Proteoform-specific interaction partners

Some interesting findings of proteoform-specific interactors are discussed in the following section.

For MAVS, the pairwise test between FL and PR resulted in 10 significant proteins (**Figure 7.A**), being PTK7, PLK1, HNRNPUL2, CEP55, CD2AP, ARRDC1, DAG1, EIF3K, TRAF2 and NUP205. Of these, only TRAF2 and PLK1 have been reported above as candidate interaction partners for MAVS FL and PR respectively, and are thus withheld as proteins that possibly interact differently with the MAVS proteoforms. By this stringent filtering, we thus remove several proteins that, although they seem to interact differently with MAVS FL or PR, they are unlikely to be interaction partners. In fact, this is obvious by visualizing their intensities profiles in the different samples before imputation (**Figure 7.B**), which shows that TRAF2 and PLK1 interact with MAVS with a significant difference in intensity between the FL and PR interactomes. For comparison, we also show the profile of two proteins that were reported as significantly different between the FL and PR interactomes, but were removed as these were not listed as candidate interactors. A first example is DAG1, which was not identified in any of the MAVS interactomes, while the found difference in the interactomes of the MAVS proteoforms is only due to differences in the imputed intensity values. In fact, this example highlights an imputation-based shortcoming of the data analysis software however, imputation is necessary for statistical analysis. A second example is ARRDC1, which is identified in almost all samples with an apparent lower intensity in the MAVS samples, making it thus unlikely that ARRDC1 is an interaction partner of MAVS. To conclude, it seems that MAVS PR has lost the interaction with TRAF2, which could be due to the fact that in MAVS PR, the domain required for interaction with TRAF2 becomes outer N-terminal (**Figure 5.A**), which affects the interaction with TRAF2. On the other hand, MAVS PR seems to interact better with PLK1, suggesting that MAVS proteoforms can both gain and lose interactors.

**Figure 7:**
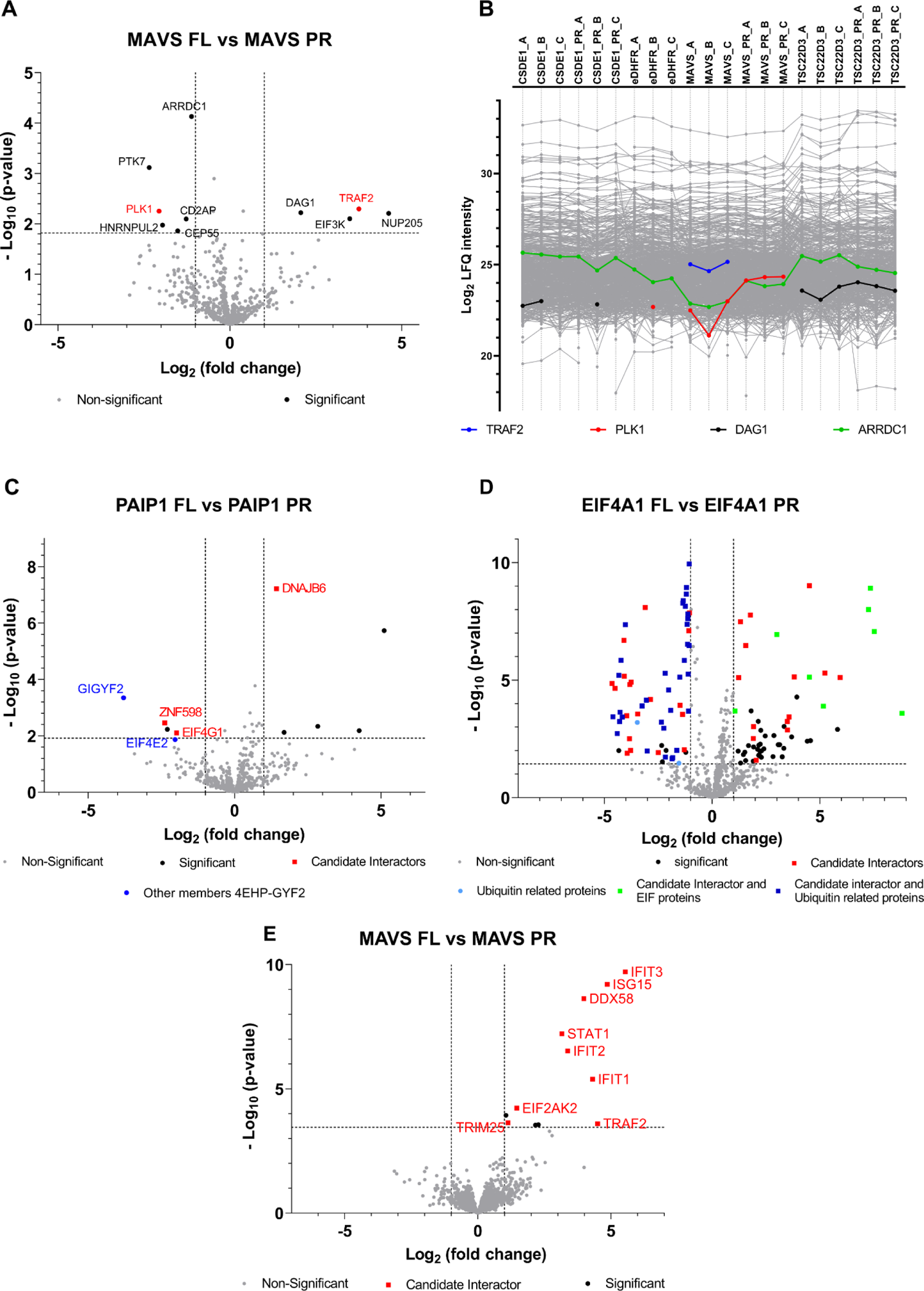
Interactomics results of selected baits with Virotrap and AP-MS. A) Volcano plot showing the candidate interaction partners that differ between MAVS FL (right) and PR (left) as identified with Virotrap. The x-axis shows the log_2_ fold change of the interactors’ intensities in the MAVS FL samples (right) relative to the MAVS PR samples (left). The y-axis shows the -log_10_ p-value of the adjusted p-value. Significant differences were defined through pairwise comparisons between MAVS FL and PR samples, with an FDR ≤ 0.05 (the corresponding –log_10_ of the adjusted p-value to a FDR of 0.05 is shown as cut-off) and a fold change larger than │1│ (cut-offs also shown on volcano plot). Proteins retained after filtering for candidate interaction partners are highlighted in red. B) Profile plot showing the label-free quantification (LFQ) intensities of all identified proteins using Virotrap. LFQ intensities of proteins quantified in every biological replicate are shown before imputation of missing values. Profiles of selected proteins reported as significantly different between MAVS FL and PR interactomes are highlighted. C) Volcano plot, similar as described in A) for PAIP1 FL and PR interactomes identified with Virotrap. Candidate interaction partners that significantly differ between PAIP1 FL and PR (FDR 0.05 and fold change > │1│log_2_ are shown in red. Other members, not listed as candidate interactors, of the 4EHP-GYF2 complex are shown in blue. D) Volcano plot similar as described in A) for EIF4A1 FL and PR interactomes as identified with Virotrap. E) Volcano plot similar as described in A) for MAVS FL and PR interactomes as identified with AP-MS.

For the polyadenylate-binding protein-interacting protein 1 (PAIP1), an N-terminal proteoform starting at position 113 (known UniProt isoform) was detected. This PAIP1 Nt-proteoform specifically interacts with GIGYF2, ZNF598 and EIF4E2, which together form the 4EHP-GYF2 complex. GIGYF2 and ZNF598 were identified from the comparison of the FL and the PR interactomes, while ZNF598 and EIF4E2 were listed as candidate interactors of PAIP1 PR (**Figure 7.C**). The engagement of PAIP1 PR with the 4EHP-GYF2 protein complex could point to a different functionality of the PAIP1 Nt-proteoform versus full-length PAIP1.

Opposite to PAIP1 PR, we report that the N-terminal proteoform (missing amino acid 1-211) of the eukaryotic initiation factor 4A-I (EIF4A1) loses several known interactions (see **Figure 7.D**). In the pairwise comparison between FL and PR, we found that the interaction with eight candidate eukaryotic initiation factors (EIF4A3, EIF4E, EIF4G3, EIF4B, EIF4G2, EIF4A2, EIF4H and EIF4G1), seems to be specific for the FL, while on the side of the EIF4A1 proteoform, we identified amongst the significant candidate interaction partners several proteins involved in proteasome-mediated protein degradation.

### 5. Y2H screens to validate the interaction profile of N-terminal proteoforms

To support our previous findings on the interactome of N-terminal proteoforms, we performed a Y2H screen similar as reported in [40, 61]. Yeast two-hybrid (Y2H) screens were performed in which all baits (both canonical protein and N-terminal proteoform), fused to the Gal4 DNA binding domain (DB), were tested against proteins encoded by the hORFeome v9.1 collection containing 17,408 ORF clones fused to the Gal4 activation domain (AD). Following first-pass screening, each bait was pairwise tested for interaction with all the candidate partners identified for any proteoform of that gene. Pairs showing a positive result were subjected to a pairwise retest and PCR products amplified from the final positive pairs were sequenced to confirm the identity of clones encoding each interacting protein. This resulted in the identification of 39 high confident binary protein-protein interactions (listed in **Supplementary Table S9**). Note that not for all baits protein-protein interactions are reported, which is due to the auto-activation of the reporter gene for some baits, while for other baits no interactions were found or did not result in a positive pair after pairwise tests.

Out of the 39 high confidence binary PPI’s reported with Y2H, 11 (28.2 %) are also reported as candidate interaction partners by Virotrap (indicated in **Supplementary Table S9**). As in general the overlap between different PPI methods is not so high [66–68], this relative high overlap shows the quality of both datasets. As an example, for EIF4A1, the specific interaction of the canonical protein (and not the PR) with PDCD4 (a well-known interaction partner), is supported by both Virotrap and Y2H.

### 6. Studying selected differences between proteoform interactomes by AP-MS

We selected three baits, MAVS, EIF4A1 and PAIP1, for further validation by AP-MS. Both FL- and PR-bait-FLAG fusion constructs were generated in which FLAG is fused to the C-terminus of the bait to avoid steric hindrance of the tag on the bait’s N-terminus. Four biological repeats of pull-down experiments using FLAG-tagged baits and an eDHFR-FLAG control were performed. Quantitative mass spectrometry was used to quantify the interaction partners of all proteoforms. In total, 1,903 proteins were identified over all experiments (**Supplementary Table S10**). Pairwise contrasts between control-bait samples and between proteoform samples were selected at a Benjamini–Hochberg adjusted *p*-value (FDR) ≤ 0.05 (**Supplementary Table S10**, second tab). Such tests between eDHFR control samples and baits resulted in 14 candidate interaction partners for MAVS FL, while for MAVS PR, only the bait was found as being significant (see **Table 4** and **Supplementary Table S10**, second tab).

**Table 4:**
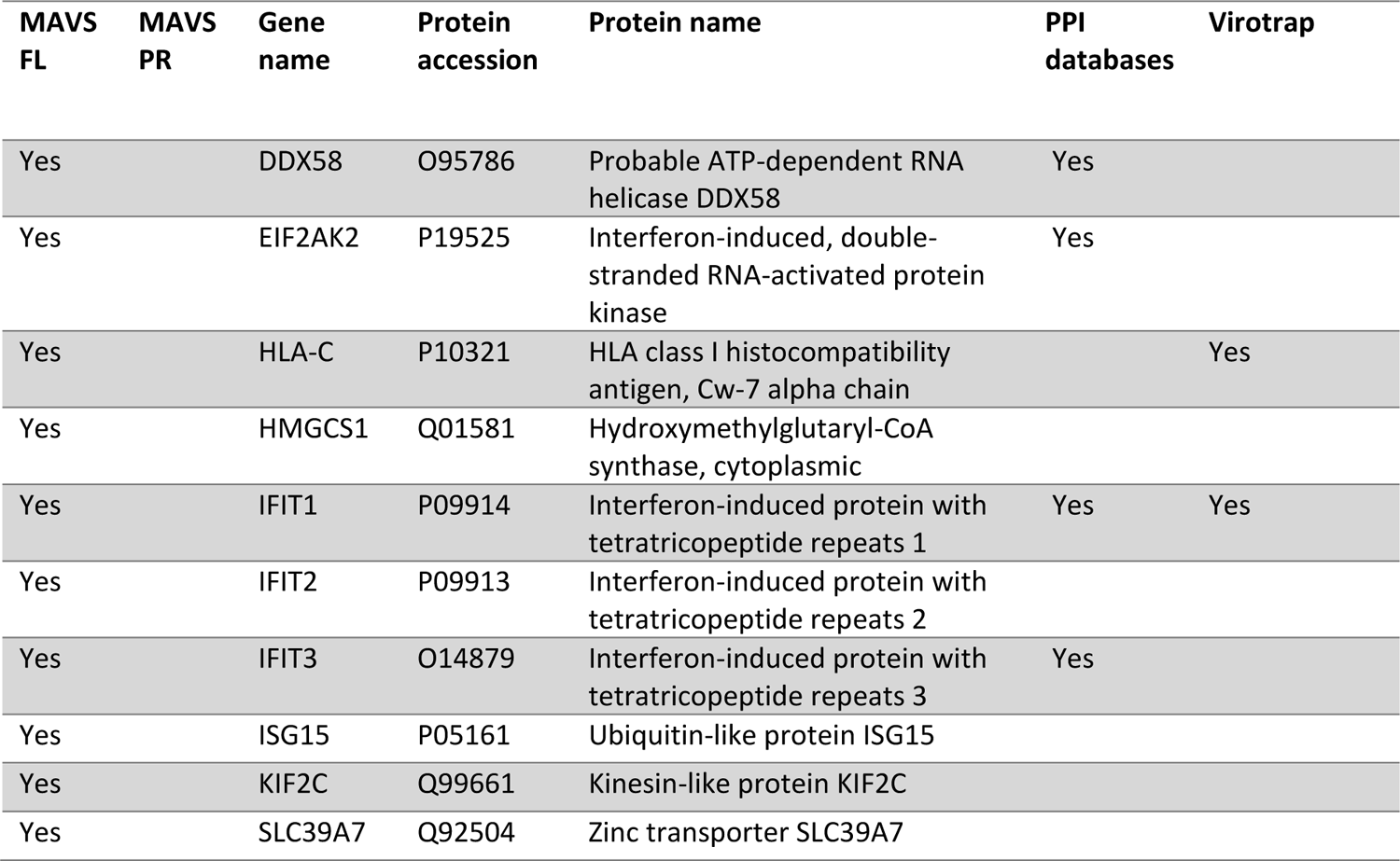
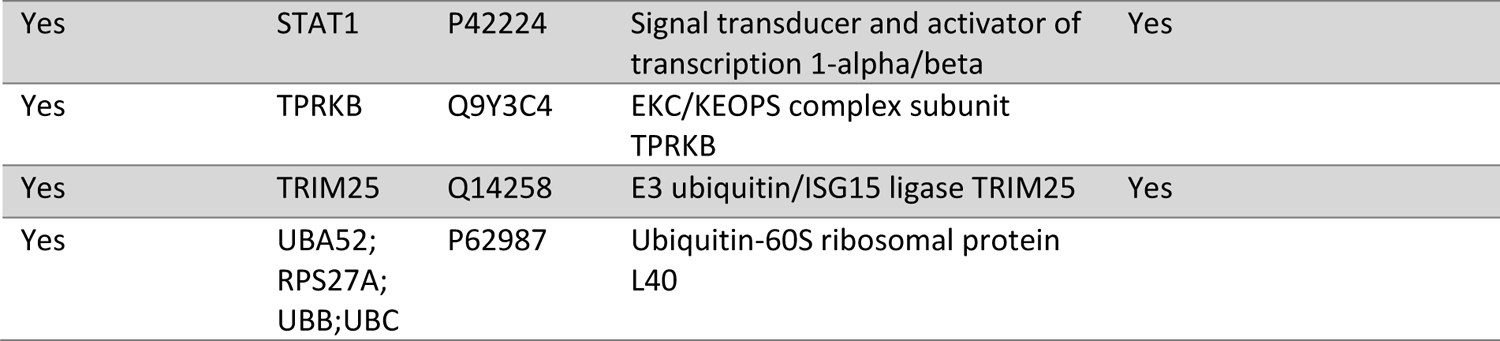
List of MAVS candidate interaction partners identified by AP-MS. MAVS AP-MS experiments, with FL or PR bait, were performed with four replicates and were challenged against eDHFR control samples. The first two columns indicate if a protein was identified as a candidate MAVS interaction partner for the FL or PR. The last but one column shows if a protein is a known MAVS interaction partner listed in BioGRID, STRING and/or IntAct. The last column indicates the overlap with the candidate interaction partners identified by Virotrap.

We compared our list of potential interaction partners with known interactors listed in BioGRID [51], STRING [38] and IntAct [60], and with candidate interactors identified by Virotrap. Six out of 14 candidate interaction partners were reported in at least one of the consulted databases, while just two candidates were also reported by Virotrap. The rather high overlap with known interaction partners points to the quality of the AP-MS data, while the overlap with the Virotrap results is a bit lower.

When applying the same selection criteria as used in the pairwise tests between Virotrap data for FL and PR, the pairwise comparison between the MAVS FL and PR interactomes reveals several candidate interaction partners that are enriched in the MAVS FL interactome (**Figure 7.E** and **Supplementary Table S10**, tab 4). In fact, our AP-MS data support our Virotrap findings that TRAF2 is enriched in MAVS FL interactomes and thus that this interaction is affected for the Nt-proteoform. Virotrap also reported the specific interaction of MAVS PR with PLK1 however, PLK1 was not identified in our AP-MS study.

For PAIP1, we identified several candidate interactors of both FL and PR, nine and 12 respectively, of which seven are known interaction partners and two were also reported as interaction partners by Virotrap (**Supplementary Table S10**). However, none of the members of the 4EHP-GYF2 complex were found as candidate interactions partners or as different between the FL and PR interactomes. We could thus not support these specific Virotrap findings by AP-MS. We hypothesize that this could be due to the differences in the PPI techniques as Virotrap allows the detection of weaker and transient interactions due to the avidity effect of multiple bait copies lining the inside of the VLP. The pairwise comparison between PAIP1 FL and PR resulted in one significant protein, UBE2T, which is reported to be enriched in proteoform samples. However, this protein was not listed as a candidate interaction partner before.

For EIF4A1 FL, only two candidate interactors were found; EIF4A2 (known interaction partner, also reported by Virotrap) and IFNA2. For the corresponding PR, no candidate interactors could be identified. These two proteins were also reported as significant in the comparison between the FL and PR, with the proteins found to be specific for the FL. Our AP-MS data thus supports that the interaction of EIF4A1 with EIF4A2 is lost for the N-terminal proteoform, which was also reported by Virotrap.

Based on our Virotrap and AP-MS interactomics data for both the canonical protein and N-terminal proteoform of MAVS and cross-checked with known interactors listed in BioGRID [51], STRING [38] and IntAct [60], we generated a protein-protein interaction network of MAVS (**Figure 8**), showing all identified candidate interaction partners. Each edge represent an identified interaction between the bait (either MAVS FL or PR, red nodes) and prey protein. In total, we identified 65 proteins and 115 interactions. The majority of the interaction partners (38) are shared between the canonical protein and the N-terminal proteoform (clustered in the middle). However, for both MAVS FL and PR, we also report a set of unique interaction partners (clustered on the left and right side). For the canonical protein, we found 16 preys that to solely interact with the canonical protein, while for the proteoform we report nine unique interaction partners.

**Figure 8:**
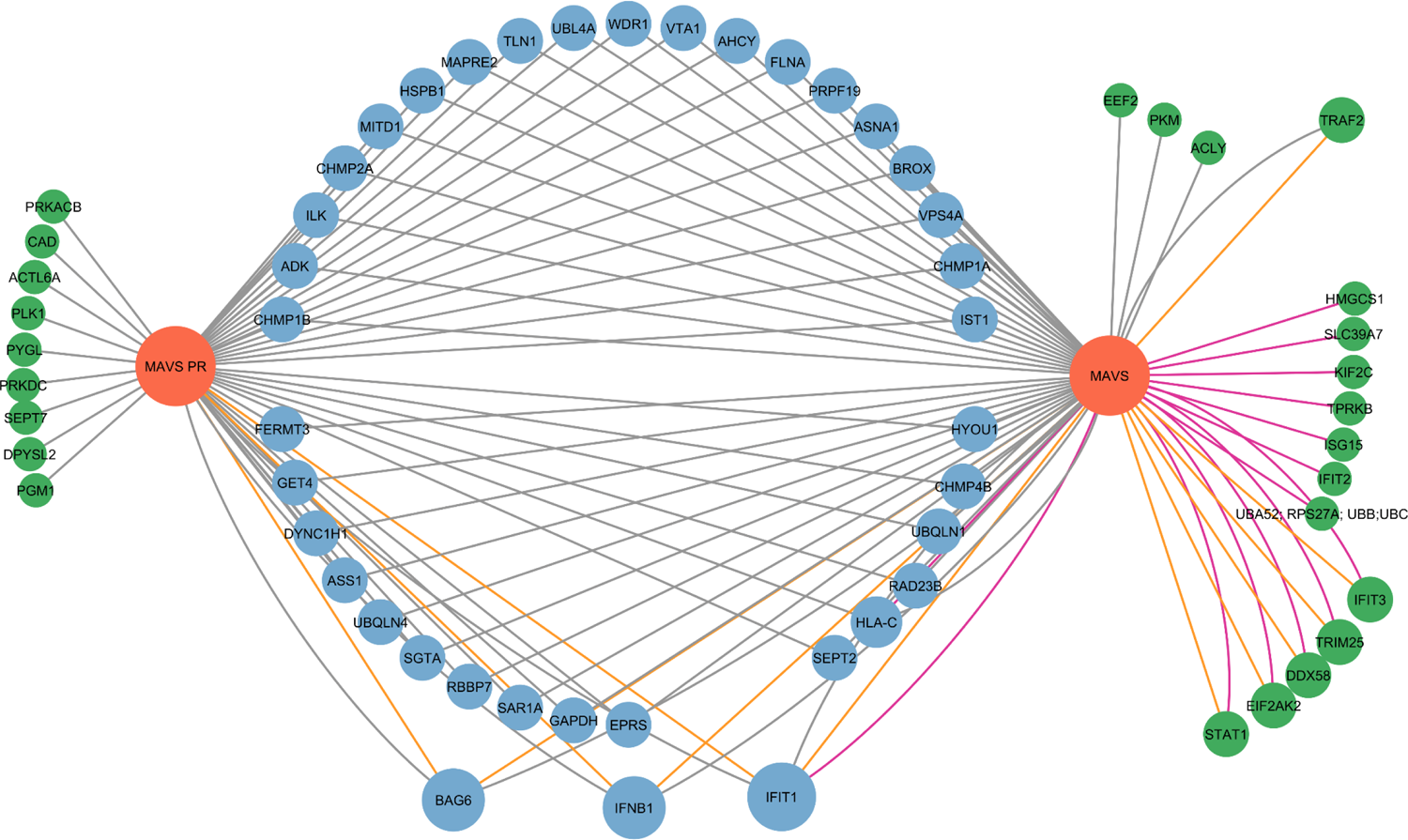
Protein-protein interaction network for proteoforms of MAVS. This network was generated by cytoscape (version 3.9.1). Bait proteins (MAVS FL and MAVS PR) are shown in red. All nodes represent interaction partners identified by either Virotrap and/or AP-MS. Grey edges indicate interactions identified by Virotrap, while purple edges indicate interactions reported by AP-MS. Orange edges represent that the interaction, reported by either Virotrap or AP-MS is supported by public PPI databases (either BioGrid, STRING or IntAct). The size of the nodes is related with how many edges are linked to this node, indicative of higher confidence for a given prey. In total, the network contains 65 nodes (proteins) and 115 edges (interactions). The interactions shared between the canonical protein and the N-terminal proteoform are clustered in the middle (blue nodes), while the proteins on the left and right side represent unique interaction partners of the N-terminal proteoform and canonical protein respectively (green nodes).

## Discussion

Several studies have reported on the chemical diversity of N-terminal proteoforms [9, 11, 23–26, 28-33] however, their functional diversity has not been investigated on a large scale. We here used positional proteomics to build a map of N-terminal proteoforms in the HEK293T cellular cytosol and identified 1,044 N-terminal proteoforms (**Table 1)**. From this map, 20 pairs of N-terminal proteoforms and their canonical protein were selected to map their protein-protein interactions (**Figure 4, Table 2**). Interaction networks of all proteins were generated, as their quality was validated by checking the overlap with known interactors listed in BioGRID [50], STRING [51] and IntAct [52] (**Supplementary Table S1, S7 and S8**). On average, N-terminal proteoforms share >60 % of their interactions with their canonical counterpart (**Figure 5D, Supplementary Table S6, second tab**). However, for all studied pairs, we could report interactome differences, suggesting functional divergence between proteoforms and noticed both the loss of interactions as well as the engagement of proteoforms in novel interactions (**Figure 8**).

Upon analyzing data of all pairs, some proteins were found as candidate interaction partners for several baits, that have highly different functions. Therefore, similar as described in [44, 69], one could think of challenging potential interactors with a list of all identified interactors in different, unrelated Virotrap experiments. In this way, proteins found in a large fraction of the whole dataset may be proteins present in VLPs irrespective of the bait and could be removed [70]. We compiled a list of all identified proteins (and their frequency) for all FL-PR (and PR1-PR2) comparisons and an alike list of all candidate interaction partners found in the bait-control comparisons (**Supplementary Table S11)**. Of note, CHMP4B and RAD23B are listed as candidate interactors of 17 baits and are thus more likely not true interactors. Among the frequently identified candidate interaction partners we also find GAG (in 11 out of 20 comparisons). This is not unexpected as we reported differences in intensities between eDHFR-GAG levels and bait-GAG levels (**Supplementary Figure S3**). As a consequence, we also found several known GAG interaction partners among the frequently identified candidate interactors such as BROX (reported for 14 baits), SLC3A2 and TSG101 (12 baits) and PDCD6IP (11 baits). For the comparisons between proteoforms (24 in total), we identified a total of 416 proteins that differ between at least one pair of FL/PR. MLF2 is reported in 11 comparisons. These are unlikely to point to specific differences between FL and PR interactomes, but are again more likely contaminants. Many proteasome-related proteins were frequently identified, as shown above for EIF4A1 (**Figure 7.D**), which indicates that one should also consider the possibility that such proteins might not be true interactors of a bait but rather inherently present in Virotrap VLPs. Of note, using AP-MS, we did not identify such proteins as candidate interaction partners for any of our baits, which strengthens our hypothesis that such proteins might be contaminants inherent to the Virotrap method.

Validation of Virotrap data was attempted by Y2H on all 20 pairs of proteoforms. However, as only a total of 39 PPIs was found by Y2H, we could not provide a lot extra evidence supporting our Virotrap findings. This low number of PPIs found by Y2H could be explained by the technology’s limitations. Y2H was reported to not work for a majority of baits due to auto-activation or failure of nuclear localization of bait and/or prey, and typically reports strong PPIs [40, 71]. Nevertheless, we were able to validate the interaction of PDCD4 with the full-length EIF4A1 protein, reported by Virotrap.

For MAVS, PAIP1 and EIF4A1, AP-MS experiments were performed to validate interactome differences between the FL and PR reported by Virotrap. For the N-terminal proteoform of MAVS, we could validate the loss of interaction with TRAF2, while for the N-terminal proteoform of EIF4A1 the loss of interaction with EIF4A2 was validated. Note that none of the other EIF proteins that Virotrap reported as specific interaction partners of the canonical EIF4A1 protein nor the interaction of the N-terminal EIF4A1 proteoform with proteasome and ubiquitin-related proteins was validated by AP-MS. For PAIP1 we were not able to validate the interaction of its N-terminal proteoform with members of the 4EHP-GYF2 complex. The differences between the candidate interactors reported with Virotrap or AP-MS might be inherently caused by the methods used. In AP-MS, samples are lysed, which likely leads to mixing of proteins that are not in the same localization, leading to false positive PPIs. In addition, several washing steps are needed, which break weaker and transient interactions. Virotrap on the other hand, traps the protein complexes in VLPs, protecting them during purification. Moreover, the GAG-grid like structure creates an avidity effect for the bait and thus in general, Virotrap allows to detect weaker and transient interactions [44, 72]. As both methods detected known protein-protein interactions, one cannot question the quality of both datasets. The fact that both methods also identified partially different interactomes for FL and PR baits, proves our working hypothesis and moreover, once again shows the complementarity of different PPI methods.

Other interesting differences were also evident from the AP-MS data (**Figure 7.E**). For instance, it was reported that IFIT3 interacts with MAVS through the N-terminus of IFIT3 [73]. However, as this interaction is enriched in the MAVS FL interactome, it hints to the fact that the N-terminus of MAVS is also important for this interaction. Besides IFIT3, several other (related) proteins are also listed to differ between FL and PR (being enriched as FL binders), and these proteins include IFIT1, IFIT2, IFIT5, ISG15 and STAT1. IFIT3 is known to interact with these proteins (see interaction network of IFIT3 as listed by STRING [38]) and IFIT1, IFIT2 and IFIT3 form a protein complex [74]). Upon viral infection, IFIT3 expression is upregulated which limits the replication of RNA viruses (by direct inhibition of translation) however, IFIT3 has also an indirect antiviral effect through its interaction with MAVS. When interferon (IFN) signaling is activated, IFIT3 induces expression of IFIT1 and IFIT2, which form a complex and stimulate TBK1 phosphorylation through MAVS. This leads to IRF3 phosphorylation and IFNβ gene expression, resulting in the upregulation of several ISGs and the phosphorylation STAT1 [75], activating canonical IFN signaling. Our interactome data lets us hypothesize that IFIT3 interacts with MAVS through the N-terminus of MAVS and that several other proteins listed as significant in the comparison between the MAVS FL and PR interactomes are due to their involvement in the IFN signaling pathway in which the interaction between IFIT3 and MAVS plays a central role. It has been reported that the N-terminal truncated proteoform of MAVS we have identified, becomes the dominant expressed MAVS proteoform upon viral infection. Moreover, this N-terminal proteoform reduces antiviral responses by interfering with interferon production and STAT1 phosphorylation [76].

In this study, Virotrap was used to map protein-protein interactions. However, as this method currently works best in HEK293T cells on cytosolic proteins, this limits the N-terminal proteoforms that can be studied. The localization of various N-terminal proteoforms is known to be affected [8, 14–22], making it possible that N-terminal proteoforms exert similar or different functions at different localizations. However, this interesting possibility could not be investigated as Virotrap is mainly restricted towards cytosolic proteins, but other approaches such as proximity labeling could be used in the future for this type of analysis. Moreover, in Virotrap, the N-terminus of the bait is fused to the C-terminus of GAG (the N-terminus of GAG is essential for its coupling to the plasma membrane). The N-terminus of the bait is thus not free and the neighborhood of GAG might sterically hinder prey proteins from interacting with the N-terminus of the bait.A decoupled variant of Virotrap, where GAG and bait are free and only coupled together upon addition of a dimerizer, could avoid this issue.

Our inspection of publicly available tissue data from different tissues [55] showed that N-terminal proteoforms are expressed in different tissues, increasing their biological relevance, but also showed that some proteoforms are expressed in a tissue-specific manner. We only mapped PPI interactions in HEK293T cells, but it seems interesting to study PPIs of N-terminal proteoforms in these tissues (or conditions, e.g. stress conditions) where these are normally expressed in.

In summary, we report confident maps of PPIs of 20 pairs of N-terminal proteoform(s) and their canonical proteins that can be explored further by the research community. Overall, our results show that N-terminal proteoforms expand the functional diversity of a proteome, which highlights the importance of considering proteoforms when studying the function of a given protein. Moving forward, studies mapping the proteoforms and their interactors in different tissues and conditions would help our understanding of the functional complexity of the proteome and our understanding of disease pathologies. However, performing such studies in a more systematic way remains challenging and labor intensive. Further improvements in PPI methods that would allow large-scale screens seem necessary.

## Acknowledgement

Contributions: K.G. conceived and supervised the project. A.B. and T.V.d.S. performed all molecular cloning. A.B and A.S. designed and performed the COFRADIC experiments. D.F. generated the custom database. A.B. and D.F. designed the filtering strategy and generated the catalogue of N-terminal proteoforms. D.F. further annotated the generated database. A.B. and K.G designed the selection priorities and selected the final pairs. C.S. extracted the data from the human protein atlas. A.B., T.V.d.S. and S.E. designed the Virotrap experments. A.B. and T.V.d.S. performed the Virotrap experiments. M.V. performed the statisitcal analysis of the Virotrap data. A.B. analyzed the Virotrap data. K.S, T.H. and M.A.C. designed and performed the Y2H experiments. A.B. and T.V.d.S performed the AP-MS experiments. A.B. and D.F. performed LC-MS/MS analysis. D.F. reannalyzed the Pandey dataset to see tissue expression of N-terminal proteoforms. A.B. and D.F. wrote the original draft of the manuscript. A.B., D.F. and K.G. edited the original draft of the manuscript. All authors edited and contributed to the final manuscript.

Acknowledgements:

This work was supported by The Research Foundation—Flanders (FWO), project numbers G008018N and G002721N (to K. G.). The authors would like to thank Katie Boucher, Evy Timmerman and Francis Impens from the VIB Proteomics Core (VIB-UGent) for operating the MS instrument.

## Data availability

The mass spectrometry proteomics data have been deposited to the ProteomeXchange Consortium via the PRIDE [77] partner repository with the following dataset identifiers:

- PXD030601 (cytosolic N-terminal COFRADIC data from the three different proteases, searched with the custom database)
- PXD039392 (“N-terminal COFRADIC on cytosolic proteins of HEK293T cells - UniProt search), data is private before publishing and can be accessed with the following login credentials: username: and password: qitZKt8a
- PXD039127 (mapping the cytosolic proteins of HEK293T cells), data is private before publishing and can be accessed with the following login credentials: username: and password: BRxGRPiR
- PXD039171 (Virotrap-based interactome analysis of 20 pairs of N-terminal proteoform(s) and their canonical counterpart), data is private before publishing and can be accessed with the following login credentials: username: and password: XRi3kNxi
- PXD039085 (AP-MS data), data is private before publishing and can be accessed with the following login credentials: username: and password: JszQHgfg
- PXD039339 (Tissue expression of N-terminal proteoforms, re-analysis “Draft map of the human proteome”), data is private before publishing and can be accessed with the following login credentials: username: and password: ob2qzRe9

## References

1. Smith, L.M., N.L. Kelleher, and P. Consortium for Top Down, Proteoform: a single term describing protein complexity. Nat Methods, 2013. 10(3): p. 186–7.

2. Bogaert, A., E. Fernandez, and K. Gevaert, N-Terminal Proteoforms in Human Disease. Trends Biochem Sci, 2020. 45(4): p. 308–320.

3. Gawron, D., K. Gevaert, and P. Van Damme, The proteome under translational control. Proteomics, 2014. 14(23-24): p. 2647–62.

4. Aebersold, R., et al., How many human proteoforms are there? Nat Chem Biol, 2018. 14(3): p. 206–214.

5. Van Damme, P., et al., N-terminal proteomics and ribosome profiling provide a comprehensive view of the alternative translation initiation landscape in mice and men. Mol Cell Proteomics, 2014. 13(5): p. 1245–61.

6. Menschaert, G., et al., Deep proteome coverage based on ribosome profiling aids mass spectrometry-based protein and peptide discovery and provides evidence of alternative translation products and near-cognate translation initiation events. Mol Cell Proteomics, 2013. 12(7): p. 1780–90.

7. Helsens, K., et al., Bioinformatics analysis of a Saccharomyces cerevisiae N-terminal proteome provides evidence of alternative translation initiation and post-translational N-terminal acetylation. J Proteome Res, 2011. 10(8): p. 3578–89.

8. Kazak, L., et al., Alternative translation initiation augments the human mitochondrial proteome. Nucleic Acids Res, 2013. 41(4): p. 2354–69.

9. Gawron, D., et al., Positional proteomics reveals differences in N-terminal proteoform stability. Mol Syst Biol, 2016. 12(2): p. 858.

10. Goyama, S., J. Schibler, and J.C. Mulloy, Alternative translation initiation generates the N-terminal truncated form of RUNX1 that retains hematopoietic activity. Exp Hematol, 2019. 72: p. 27–35.

11. Claus, P., et al., Differential intranuclear localization of fibroblast growth factor-2 isoforms and specific interaction with the survival of motoneuron protein. J Biol Chem, 2003. 278(1): p. 479–85.

12. Hartmann, E.M. and J. Armengaud, N-terminomics and proteogenomics, getting off to a good start. Proteomics, 2014. 14(23-24): p. 2637–46.

13. Varshavsky, A., N-degron and C-degron pathways of protein degradation. Proc Natl Acad Sci U S A, 2019. 116(2): p. 358–366.

14. Kobayashi, R., et al., Targeted mass spectrometric analysis of N-terminally truncated isoforms generated via alternative translation initiation. FEBS Lett, 2009. 583(14): p. 2441–5.

15. Kunze, M. and J. Berger, The similarity between N-terminal targeting signals for protein import into different organelles and its evolutionary relevance. Front Physiol, 2015. 6: p. 259.

16. Rossmanith, W., Localization of human RNase Z isoforms: dual nuclear/mitochondrial targeting of the ELAC2 gene product by alternative translation initiation. PLoS One, 2011. 6(4): p. e19152.

17. Suzuki, Y., et al., An upstream open reading frame and the context of the two AUG codons affect the abundance of mitochondrial and nuclear RNase H1. Mol Cell Biol, 2010. 30(21): p. 5123–34.

18. Leissring, M.A., et al., Alternative translation initiation generates a novel isoform of insulin-degrading enzyme targeted to mitochondria. Biochem J, 2004. 383(Pt. 3): p. 439–46.

19. Land, T. and T.A. Rouault, Targeting of a human iron-sulfur cluster assembly enzyme, nifs, to different subcellular compartments is regulated through alternative AUG utilization. Mol Cell, 1998. 2(6): p. 807–15.

20. Packham, G., M. Brimmell, and J.L. Cleveland, Mammalian cells express two differently localized Bag-1 isoforms generated by alternative translation initiation. Biochem J, 1997. 328 **(Pt** **3**): p. 807–13.

21. Monteuuis, G., et al., Non-canonical translation initiation in yeast generates a cryptic pool of mitochondrial proteins. Nucleic Acids Res, 2019. 47(11): p. 5777–5791.

22. Li, Y.R. and M.J. Liu, Prevalence of alternative AUG and non-AUG translation initiators and their regulatory effects across plants. Genome Res, 2020. 30(10): p. 1418–1433.

23. Zecha, J., et al., Peptide Level Turnover Measurements Enable the Study of Proteoform Dynamics. Mol Cell Proteomics, 2018. 17(5): p. 974–992.

24. Song, K.Y., et al., Differential use of an in-frame translation initiation codon regulates human mu opioid receptor (OPRM1). Cell Mol Life Sci, 2009. 66(17): p. 2933–42.

25. Calligaris, R., et al., Alternative translation initiation site usage results in two functionally distinct forms of the GATA-1 transcription factor. Proc Natl Acad Sci U S A, 1995. 92(25): p. 11598–602.

26. Thomas, D., et al., Alternative translation initiation in rat brain yields K2P2.1 potassium channels permeable to sodium. Neuron, 2008. 58(6): p. 859–70.

27. Kwon, H., et al., Alternative translation initiation of Caveolin-2 desensitizes insulin signaling through dephosphorylation of insulin receptor by PTP1B and causes insulin resistance. Biochim Biophys Acta Mol Basis Dis, 2018. 1864(6 Pt A): p. 2169-2182.

28. Khanna-Gupta, A., et al., Up-regulation of translation eukaryotic initiation factor 4E in nucleophosmin 1 haploinsufficient cells results in changes in CCAAT enhancer-binding protein alpha activity: implications in myelodysplastic syndrome and acute myeloid leukemia. J Biol Chem, 2012. 287(39): p. 32728–37.

29. Gu, S., et al., Alternative translation initiation of human regulators of G-protein signaling-2 yields a set of functionally distinct proteins. Mol Pharmacol, 2008. 73(1): p. 1–11.

30. Malaney, P., V.N. Uversky, and V. Dave, PTEN proteoforms in biology and disease. Cell Mol Life Sci, 2017. 74(15): p. 2783–2794.

31. Fukushima, M., et al., Alternative translation initiation gives rise to two isoforms of Orai1 with distinct plasma membrane mobilities. J Cell Sci, 2012. 125(Pt 18): p. 4354–61.

32. Simkin, D., E.J. Cavanaugh, and D. Kim, Control of the single channel conductance of K2P10.1 (TREK-2) by the amino-terminus: role of alternative translation initiation. J Physiol, 2008. 586(23): p. 5651–63.

33. Trulley, P., et al., Alternative Translation Initiation Generates a Functionally Distinct Isoform of the Stress-Activated Protein Kinase MK2. Cell Rep, 2019. 27(10): p. 2859–2870 e6.

34. Scott, J.D. and T. Pawson, Cell signaling in space and time: where proteins come together and when they’re apart. Science, 2009. 326(5957): p. 1220-4.

35. Perkins, J.R., et al., Transient protein-protein interactions: structural, functional, and network properties. Structure, 2010. 18(10): p. 1233–43.

36. Oughtred, R., et al., The BioGRID interaction database: 2019 update. Nucleic Acids Res, 2019. 47(D1): p. D529–D541.

37. Szklarczyk, D., et al., STRING v11: protein-protein association networks with increased coverage, supporting functional discovery in genome-wide experimental datasets. Nucleic Acids Res, 2019. 47(D1): p. D607–D613.

38. Szklarczyk, D., et al., Correction to ‘The STRING database in 2021: customizable protein-protein networks, and functional characterization of user-uploaded gene/measurement sets’. Nucleic Acids Res, 2021. 49(18): p. 10800.

39. Ghadie, M.A., et al., Domain-based prediction of the human isoform interactome provides insights into the functional impact of alternative splicing. PLoS Comput Biol, 2017. 13(8): p. e1005717.

40. Yang, X., et al., Widespread Expansion of Protein Interaction Capabilities by Alternative Splicing. Cell, 2016. 164(4): p. 805–17.

41. Ellis, J.D., et al., Tissue-specific alternative splicing remodels protein-protein interaction networks. Mol Cell, 2012. 46(6): p. 884–92.

42. Bludau, I. and R. Aebersold, Proteomic and interactomic insights into the molecular basis of cell functional diversity. Nat Rev Mol Cell Biol, 2020. 21(6): p. 327–340.

43. Staes, A., et al., Improved recovery of proteome-informative, protein N-terminal peptides by combined fractional diagonal chromatography (COFRADIC). Proteomics, 2008. 8(7): p. 1362–70.

44. Eyckerman, S., et al., Trapping mammalian protein complexes in viral particles. Nat Commun, 2016. 7: p. 11416.

45. Bogaert, A., et al., Limited Evidence for Protein Products of Noncoding Transcripts in the HEK293T Cellular Cytosol. Mol Cell Proteomics, 2022. 21(8): p. 100264.

46. Cox, J., et al., Accurate proteome-wide label-free quantification by delayed normalization and maximal peptide ratio extraction, termed MaxLFQ. Mol Cell Proteomics, 2014. 13(9): p. 2513–26.

47. Tyanova, S., et al., The Perseus computational platform for comprehensive analysis of (prote)omics data. Nat Methods, 2016. 13(9): p. 731–40.

48. Martens, L., J. Vandekerckhove, and K. Gevaert, DBToolkit: processing protein databases for peptide-centric proteomics. Bioinformatics, 2005. 21(17): p. 3584–5.

49. Kumar, M., et al., The Eukaryotic Linear Motif resource: 2022 release. Nucleic Acids Res, 2022. 50(D1): p. D497–D508.

50. Fortelny, N., et al., Proteome TopFIND 3.0 with TopFINDer and PathFINDer: database and analysis tools for the association of protein termini to pre- and post-translational events. Nucleic Acids Res, 2015. 43(Database issue): p. D290-7.

51. Oughtred, R., et al., The BioGRID database: A comprehensive biomedical resource of curated protein, genetic, and chemical interactions. Protein Sci, 2021. 30(1): p. 187–200.

52. Amberger, J.S., et al., OMIM.org: leveraging knowledge across phenotype-gene relationships. Nucleic Acids Res, 2019. 47(D1): p. D1038–D1043.

53. Huntley, R.P., et al., The GOA database: gene Ontology annotation updates for 2015. Nucleic Acids Res, 2015. 43(Database issue): p. D1057-63.

54. Thul, P.J., et al., A subcellular map of the human proteome. Science, 2017. 356(6340).

55. Kim, M.S., et al., A draft map of the human proteome. Nature, 2014. 509(7502): p. 575-81.

56. Degroeve, S., et al., ionbot: a novel, innovative and sensitive machine learning approach to LC-MS/MS peptide identification. bioRxiv, 2022: p. 2021.07.02.450686.

57. Bouwmeester, R., et al., DeepLC can predict retention times for peptides that carry as-yet unseen modifications. Nat Methods, 2021. 18(11): p. 1363–1369.

58. Ning, Z., et al., From cells to peptides: “one-stop” integrated proteomic processing using amphipols. J Proteome Res, 2013. 12(3): p. 1512–9.

59. Szklarczyk, D., et al., The STRING database in 2021: customizable protein-protein networks, and functional characterization of user-uploaded gene/measurement sets. Nucleic Acids Res, 2021. 49(D1): p. D605–D612.

60. Orchard, S., et al., The MIntAct project--IntAct as a common curation platform for 11 molecular interaction databases. Nucleic Acids Res, 2014. 42(Database issue): p. D358-63.

61. Luck, K., et al., A reference map of the human binary protein interactome. Nature, 2020. 580(7803): p. 402-408.

62. Kaulich, P.T., et al., Multi-protease Approach for the Improved Identification and Molecular Characterization of Small Proteins and Short Open Reading Frame-Encoded Peptides. J Proteome Res, 2021. 20(5): p. 2895–2903.

63. Verbruggen, S., et al., PROTEOFORMER 2.0: Further Developments in the Ribosome Profiling-assisted Proteogenomic Hunt for New Proteoforms. Mol Cell Proteomics, 2019. 18(8 suppl 1): p. S126-S140.

64. Verbruggen, S., et al., Spectral Prediction Features as a Solution for the Search Space Size Problem in Proteogenomics. Mol Cell Proteomics, 2021. 20: p. 100076.

65. Sudhir, P.R. and C.H. Chen, Proteomics-Based Analysis of Protein Complexes in Pluripotent Stem Cells and Cancer Biology. Int J Mol Sci, 2016. 17(3): p. 432.

66. Masschaele, D., et al., High-Confidence Interactome for RNF41 Built on Multiple Orthogonal Assays. J Proteome Res, 2018. 17(4): p. 1348–1360.

67. De Bodt, S., et al., Predicting protein-protein interactions in Arabidopsis thaliana through integration of orthology, gene ontology and co-expression. BMC Genomics, 2009. 10: p. 288.

68. Uszczynska-Ratajczak, B., et al., Towards a complete map of the human long non-coding RNA transcriptome. Nat Rev Genet, 2018. 19(9): p. 535–548.

69. Meysman, P., et al., Protein complex analysis: From raw protein lists to protein interaction networks. Mass Spectrom Rev, 2017. 36(5): p. 600–614.

70. Mellacheruvu, D., et al., The CRAPome: a contaminant repository for affinity purification-mass spectrometry data. Nat Methods, 2013. 10(8): p. 730–6.

71. Lievens, S., et al., Two-hybrid and its recent adaptations. Drug Discov Today Technol, 2006. 3(3): p. 317–24.

72. Thery, F., et al., Ring finger protein 213 assembles into a sensor for ISGylated proteins with antimicrobial activity. Nat Commun, 2021. 12(1): p. 5772.

73. Liu, X.Y., et al., IFN-induced TPR protein IFIT3 potentiates antiviral signaling by bridging MAVS and TBK1. J Immunol, 2011. 187(5): p. 2559–68.

74. Pichlmair, A., et al., IFIT1 is an antiviral protein that recognizes 5’-triphosphate RNA. Nat Immunol, 2011. 12(7): p. 624–30.

75. Chikhalya, A., et al., Human IFIT3 Protein Induces Interferon Signaling and Inhibits Adenovirus Immediate Early Gene Expression. mBio, 2021. 12(6): p. e0282921.

76. Brubaker, S.W., et al., A bicistronic MAVS transcript highlights a class of truncated variants in antiviral immunity. Cell, 2014. 156(4): p. 800–11.

77. Perez-Riverol, Y., et al., The PRIDE database and related tools and resources in 2019: improving support for quantification data. Nucleic Acids Res, 2019. 47(D1): p. D442–D450.

