## Supplementary Figures for "N-terminal proteoforms may engage in different protein complexes"

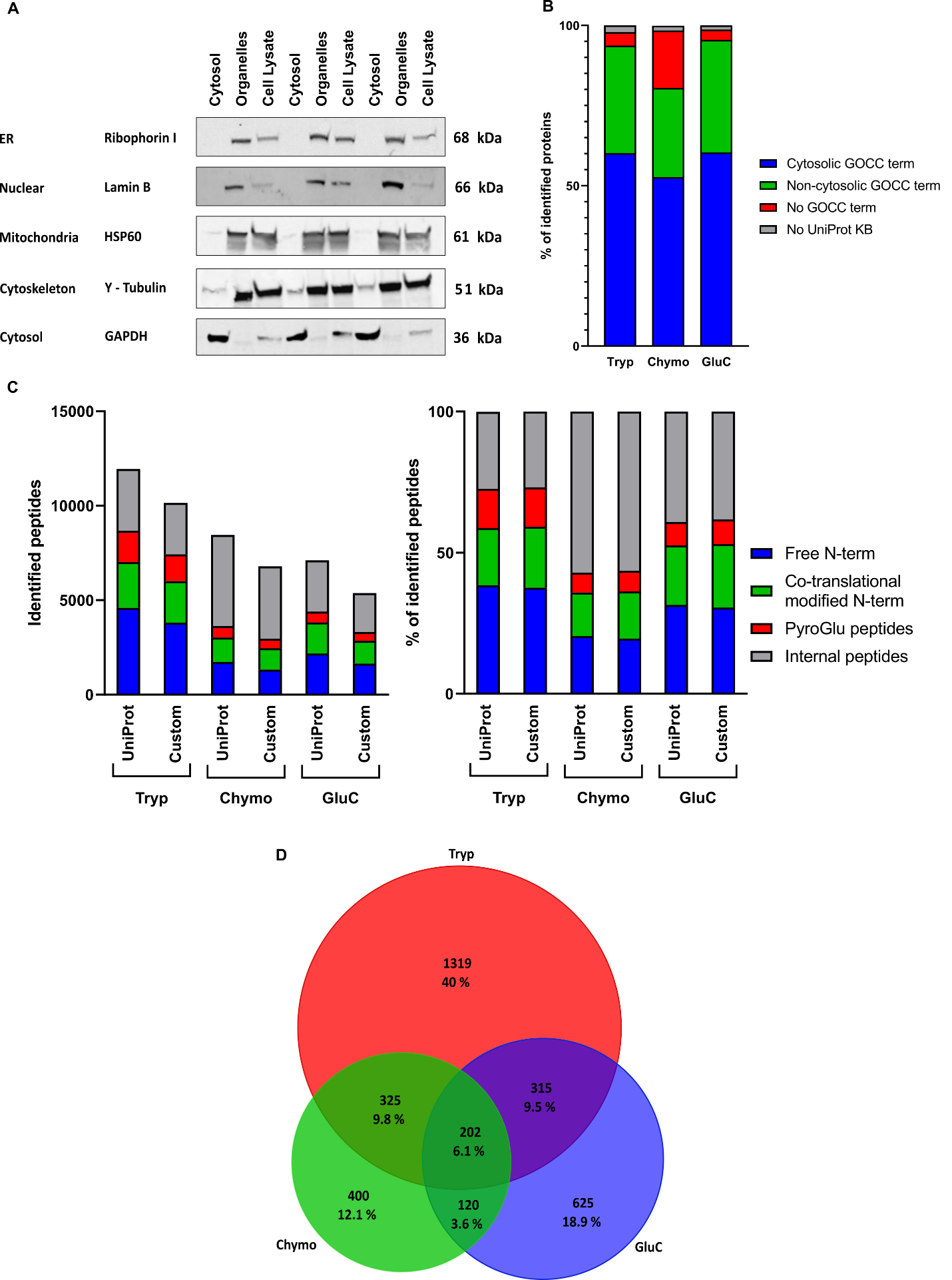

***Figure S1: Overview of the results obtained by cytosolic enrichment of HEK293T cells followed by N-terminal COFRADIC and the subsequent filtering to a catalogue of N-terminal proteoforms.*** *A) Efficiency of the use of digitonin for extracting cytosolic proteins from HEK293T cells. 10^7^ cells were collected and lysed in 1.25 ml of CFS buffer containing 0.02% digitonin. Following centrifugation, the supernatant was collected and the cell pellet was further extracted in RIPA lysis buffer. A total cell lysate was made by lysis of 10^7^ HEK293T cells in RIPA lysis buffer. An aliquot of each extract was subjected to 4-12% SDS-PAGE followed by Western blot analysis using an anti-GAPDH antibody as a marker for the cytosol, an anti-HSP60 antibody as a marker for mitochondria, an anti-Lamin B antibody as a marker for the nucleus, anti-γ-Tubulin as a marker for the cytoskeleton and an antibody against Ribophorin I as a marker for the ER. ER and nuclear markers were absent in the cytosolic fraction, whilst clearly visible in the organelle fraction and the total cell lysate. For HSP60 (mitochondrial marker) and γ-Tubulin (cytoskeleton marker), faint staining was observed in the cytosolic fraction and intense staining in the organelle fraction and the total cell lysate. GAPDH is not observed in the organelle fraction, while intense in the cytosolic fraction, and this intensity is clearly higher compared to that in the total cell lysate. B) When evaluating Gene Ontology Cellular Component terms associated with the identified proteins for “cytosol” (GO: 0005829), a comparable fraction of cytosolic proteins (around 60%) was found in the three samples. This is higher than expected when analyzing a total lysate (Fisher’s exact test,* p *< 1e-5), given that The Human Protein Atlas (Cell Atlas) reports that 4,740 proteins (24% of the human proteome) localize to the cytosol. C)* *Overview of the numbers of distinct peptides found in the different samples and N-terminal enrichment efficiencies. LC-MS/MS data were searched using Mascot and either in UniProt or our custom-build database, and all identifications were stored and retrieved via ms_lims (at a confidence interval of 0.01) [1]. Identifications were grouped by peptide sequence. When the same sequence was found several times with different N-terminal modifications, priority was given to in vivo acetylated peptides. The left panel shows the total number of peptides identified by the different proteases and their categories, while the right panel shows the percentage of each category of peptides. COFRADIC enriches for in vivo free N-terminal peptides, co-translationally modified N-terminal peptides and pyro-glutamate starting peptides and its efficiency reached to about 60% for trypsin and GluC samples and >40 % for the chymotrypsin-treated sample. D) Venn diagrams showing the overlap between the final confident N-terminal peptides identified with the different proteases. Trypsin accounts for the largest set of identified N-terminal peptides, but both GluC and chymotrypsin add an additional set of confident N-terminal peptides leading to a total increase of 1,145 N-terminal peptides.*

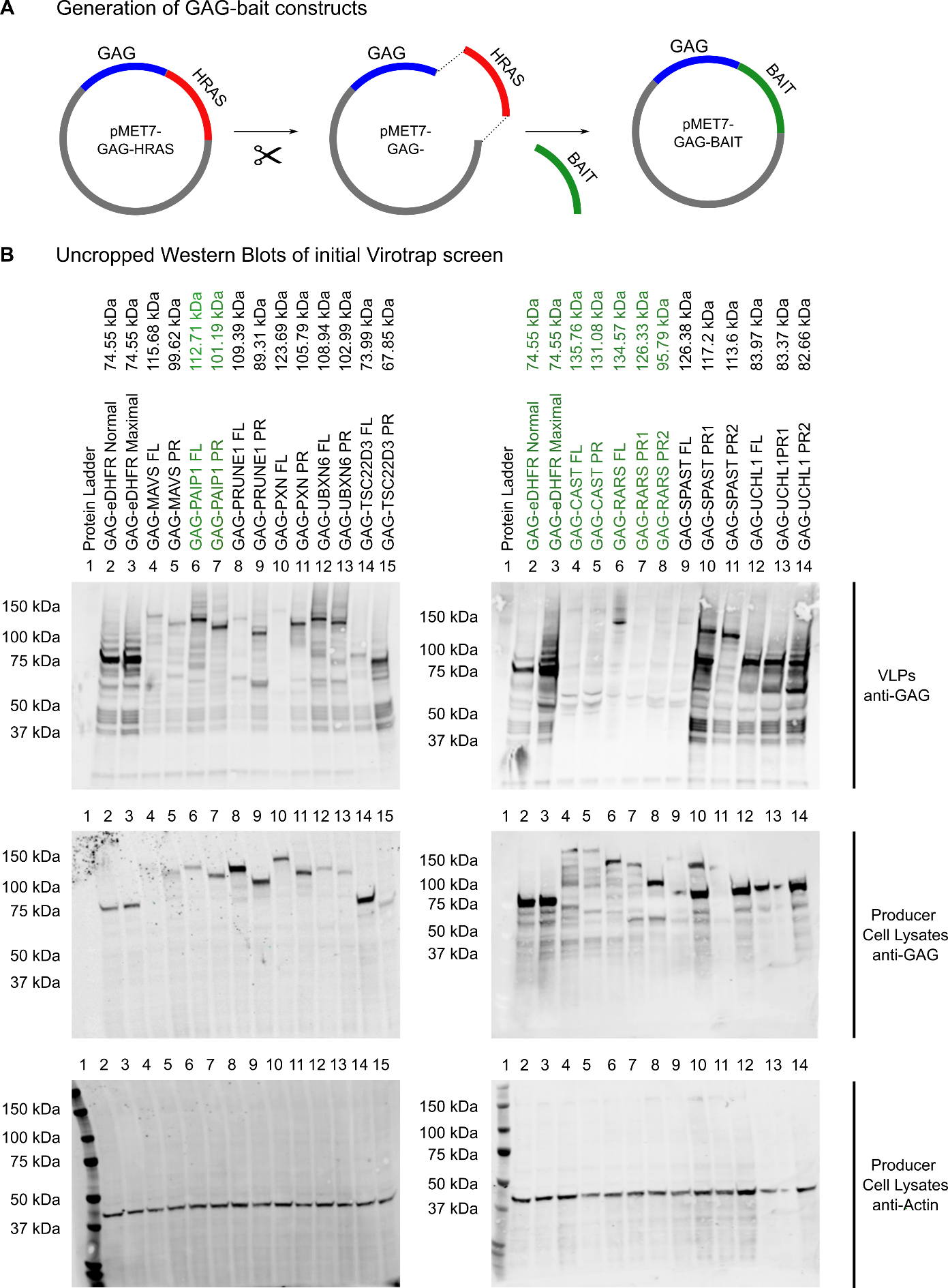

***Figure S2: Additional information Virotrap experiments.*** *A) Scheme showing the generation of plasmids containing GAG-bait. We started from the pMET7-GAG-HRAS vector from which HRAS was removed by restriction digestion. Bait sequences were amplified from cDNA or gBlocks with the required overlapping overhangs and ligated into the linearized pMET7-GAG vector with T4 DNA ligase B) Uncropped Western Blot results of Figure 3.* *Virotrap experiments for detection of expression and recruitment into the VLPs of GAG-bait fusion proteins. 1.15x10^6^ HEK293T cells were seeded and transfected after 24 h with GAG-bait constructs. Additional co-transfection of VSV-G/FLAG-VSV-G expression constructs allowed VLP purification, which was followed by direct on-bead lysis of the VLPs. The producer cells were also collected and lysed, and both lysates were subjected to 4-12% SDS-PAGE followed by Western blot using anti-GAG (bait expression levels and particles) and anti-α-actin antibodies (as loading control, 42 kDa, cell lysates). Both results of VLPs and producer cell lysates are shown. For GAG-eDHFR and GAG-PAIP1 (both FL and PR), a clear band can be detected at the desired height in both VLPs and cells. For RARS, all constructs seem well expressed in the cells but only RARS FL is recruited into VLPs. CAST is found only weakly expressed in the cell lysates (band also detected to high) and does not seem to be pulled into VLPs.*

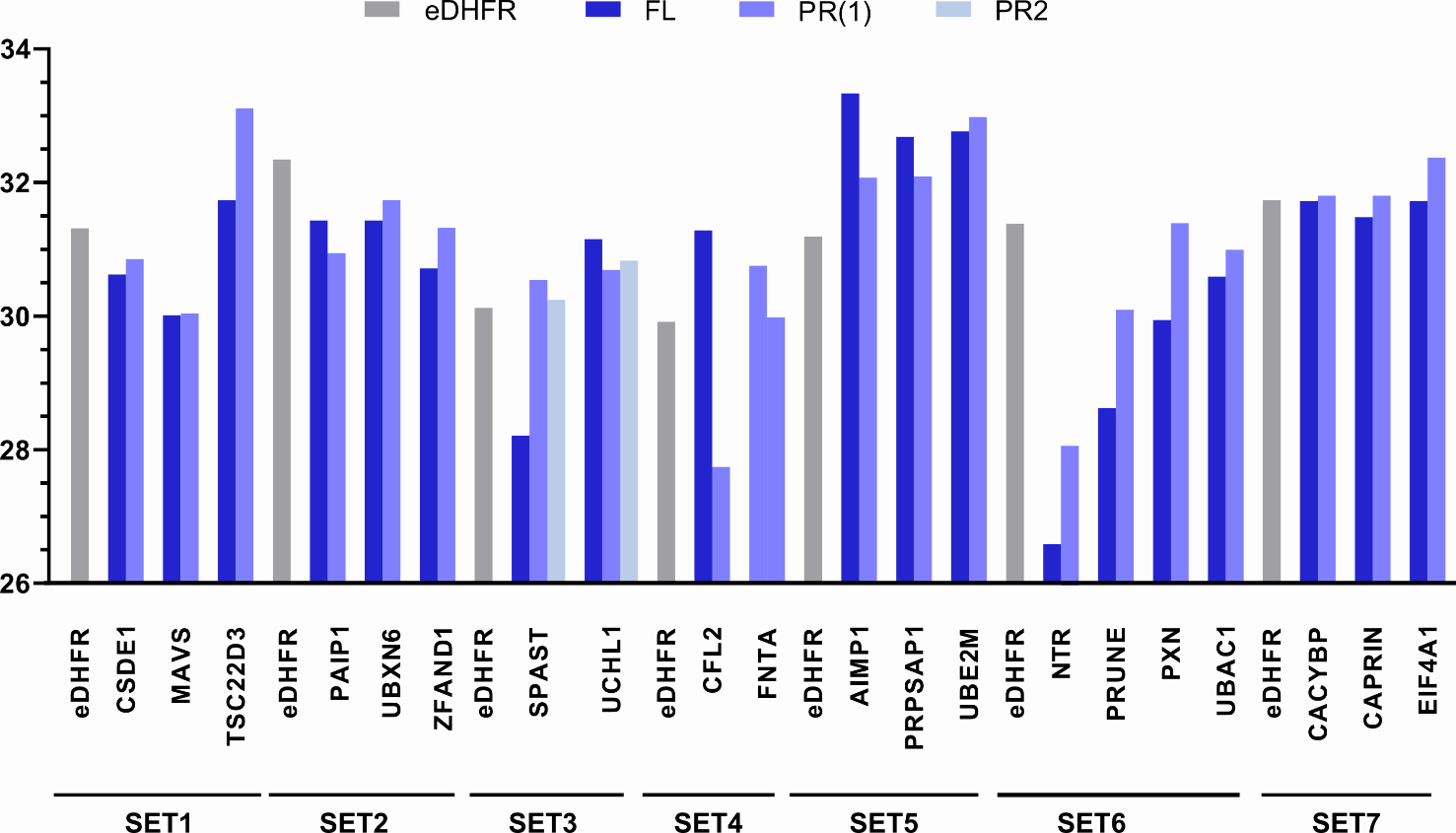

***Figure S3. Comparisons of bait intensities.*** *Bar chart showing the log_2_ transformed label free quantification (LFQ) intensities of the different baits (the average value of the three replicates is shown) used in out Virotrap experiments.*

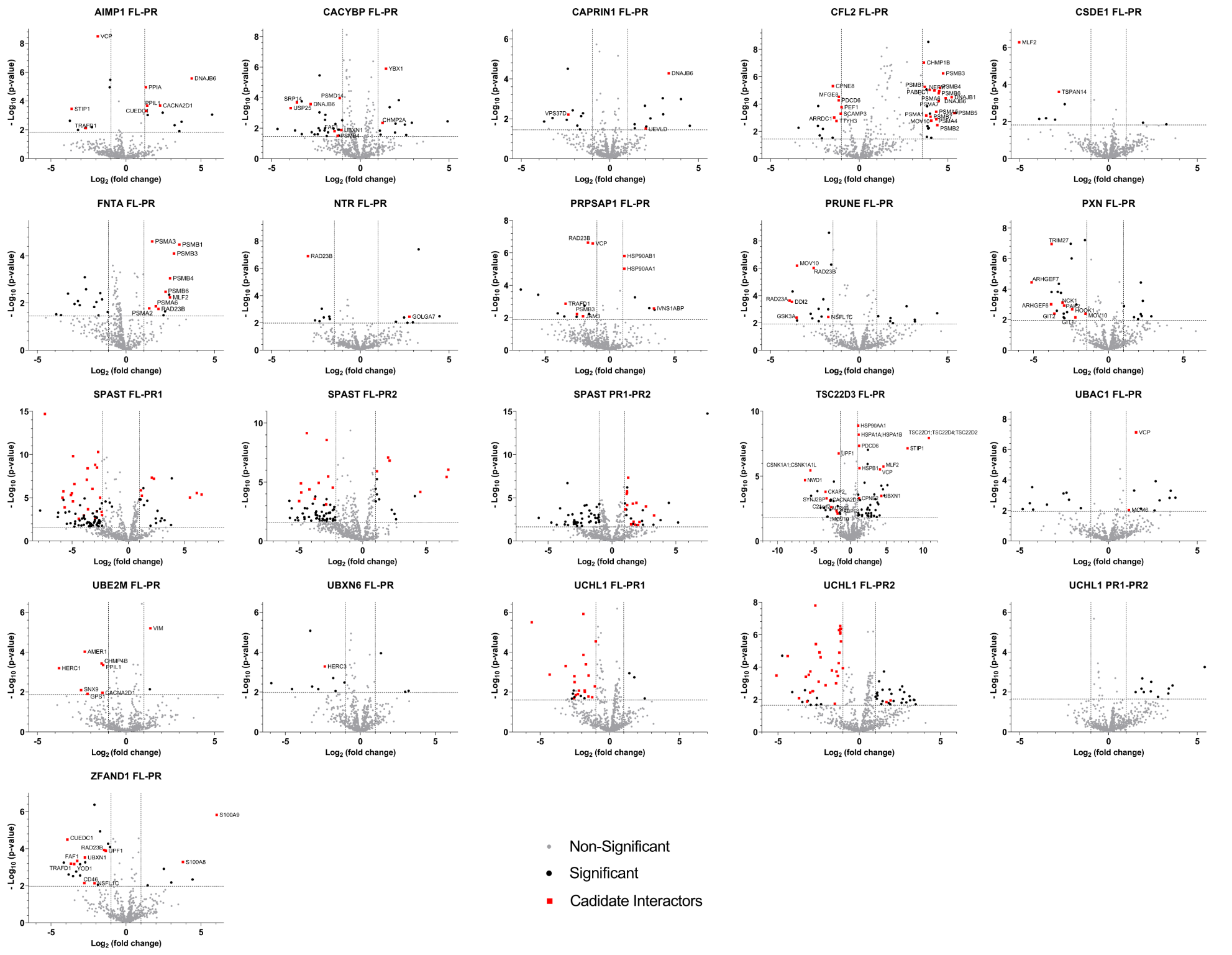

***Figure S4: Virotrap results of all pairwise teste between FL and PR of all baits.*** *Volcano plots showing the candidate interaction partners that differ between a bait FL (right) and PR (left) (or PR1 and PR2). The x-axis shows the log_2_ fold change of the interactors intensities in the bait FL samples (right) relative to the bait PR samples (left). The y-axis shows the -log_10_ p-value of the adjusted p-value. Significant differences were defined through pairwise comparisons between bait FL and PR samples, with an FDR ≤ 0.05 (the corresponding –log_10_ of the adjusted p-value to a FDR of 0.05 is shown as cut-off) and a fold change larger than │1│ (cut-offs also shown on volcano plot). Proteins retained after filtering for candidate interaction partners are highlighted in red.*

1.	Helsens, K., et al., *ms_lims, a simple yet powerful open source laboratory information management system for MS-driven proteomics.* Proteomics, 2010. **10**(6): p. 1261-4.
