## Supplementary Materials and Methods for "N-terminal proteoforms may engage in different protein complexes"

*Cytosol extraction*

Cytosolic extracts were prepared from 2.5 x10^7^ HEK293T cells as described in [1]. Cell pellets were washed with ice-cold DPBS (Thermo Fisher Scientific cat. No. 14190250) and resuspended in 1.25 ml of cell-free systems (CFS) buffer (220 mM mannitol, 170 mM sucrose, 5 mM NaCl, 5 mM MgCl_2_, 10 mM HEPES pH 7.5, 2.5 mM KH_2_PO_4_, 0.02 % digitonin and Complete protease inhibitor cocktail (Roche Cat. No. 4693132001)) and kept on ice for 2 min. Lysates were cleared by centrifugation at 14,000 x g for 15 min at 4 °C. The supernatants, containing the cytosolic extracts, were collected. The pellets and a HEK293T total lysate serving as control were resuspended in 1 ml of RIPA buffer (50 mM Tris-HCl pH 7.4, 150 mM NaCl, 1% NP40, 0.5% sodium deoxycholate, 0.1% SDS and Complete protease inhibitor cocktail) followed by three freeze-thaw cycles. Lysates were cleared by centrifugation at 16,000 x g for 15 min at room temperature (RT). 200 μl of each sample was used for Western blot analysis, while the rest of the sample was used for N-terminal peptide enrichment by COFRADIC.

*Western blot analysis*

Proteins were denatured in XT Sample buffer (Bio-Rad cat. No. 1610791) and XT reducing agent (Bio-Rad cat. No. 1610792), heated at 99 °C for 10 min and centrifuged for 5 min at 16,000 x g (room temperature). 25 μg of each protein mixture was separated by SDS-PAGE (polyacrylamide gel electrophoresis) on a Criterion XT 4-12% Bis-Tris gel (Biorad Laboratories cat. No. 3450124). Proteins were transferred to a PVDF membrane (Merck Millipore cat. No. IPFL00010) after which the membrane was blocked using Odyssey Blocking buffer (PBS) (LI-COR cat. No. 927-4000) diluted once with TBS-T (TBS supplemented with 0.1% Tween 20). Immunoblots were incubated overnight with primary antibodies against GAPDH (Abcam, ab8245), HSP60 (Santa Cruz, sc-13115), Lamin B (Santa Cruz, sc-374015), Ribophorin I (Santa Cruz, sc-12164) or γ-Tubulin (Thermo Fisher Scientific, MA1-850) in Odyssey Blocking buffer (PBS) diluted once with TBS-T. Blots were washed four times with TBS-T, incubated with fluorescent-labeled secondary antibodies (IRDye 800CW Goat Anti-Mouse IgG polyclonal 0.5 mg from LI-COR, Cat. No 926-32210) and IRDye 800CW Donkey Anti-Goat IgG polyclonal 0.5 mg from LI-COR cat. No. 926-32214) in Odyssey blocking buffer diluted once with TBS-T for 1 h. After three washes with TBS-T and an additional wash in TBS, immunoblots were imaged using the Odyssey infrared imaging system (LI-COR).

*N-terminal COFRADIC*

N-terminal peptides were enriched by COFRADIC as described previously [2] however without the pyroglutamate removal and SCX steps. In the following, we only mention the main differences with the published protocol. In brief, 1 mg of cytosolic proteins was used. As digitonin was used for cytosolic extraction in combination with the RIPA lysis buffer, which interfere with LC-MS/MS analysis, the samples were cleaned-up using Pierce® Detergent removal spin columns (Thermo Fisher Scientific, cat. No. 87777) according to the manufacturer’s instructions. Then, guanidinium hydrochloride was added to a final concentration of 4 M before proteins were reduced (with 15 mM f.c. (final concentration) TCEP) and alkylated (with 30 mM f.c. iodoacetamide) for 15 min at 37 °C. To be able to distinguish between *in vivo* Nt-acetylation events and free primary amines, all primary protein amines were blocked using stable isotope encoded acetate, i.e. an NHS-ester of ^13^C_1_D_3_-acetate. The acetylation reaction was allowed to proceed for 1 h at 30 °C and was repeated once. Prior to digestion, the samples were desalted on a NAP-10 column in 50 mM freshly prepared ammonium bicarbonate, pH 7.8. Samples were digested either with trypsin in a trypsin/substrate ratio of 1/50 (w/w) (Promega Cat. No. V5111) and incubated overnight at 37 °C, chymotrypsin in a chymotrypsin/ substrate ratio of 1/20 (w/w) (Promega Cat. No. V1061) and incubated overnight at 25 °C or endoproteinase GluC in a GluC/ substrate ratio of 1/20 (w/w) (Promega Cat. No.V1651) and incubated overnight at 37 °C. After vacuum drying, the samples were re-dissolved in 80 μl loading solvent A (2% acetonitrile (ACN), 0.1% TFA in ddH_2_O) before isolating N-terminal peptides by two subsequent RP-HPLC fractionations with a TNBS reaction in between.

*LC-MS/MS analysis of N-terminal peptides and peptide identification*

LC-MS/MS analysis was performed similar as reported before [2]. Each COFRADIC fraction was re-solubilized in 20 μl loading solvent A and half of each fraction was injected for LC-MS/MS analysis on an Ultimate 3000 RSLCnano system in-line connected to an Orbitrap Fusion Lumos mass spectrometer (Thermo Scientific, Germany). Trapping was performed at 10 μl/min for 4 min in loading solvent A on a 20 mm trapping column (made in-house, 100 μm internal diameter (I.D.), 5 μm beads, C18 Reprosil-HD, Dr. Maisch, Germany). The peptides were separated on a 200 cm μPAC™ column (C18-endcapped functionality, 300 μm wide channels, 5 μm porous-shell pillars, inter pillar distance of 2.5 μm and a depth of 20 μm; Pharmafluidics, Belgium). The column was kept at a constant temperature of 50 °C. Peptides were eluted by a linear gradient reaching 33% MS solvent B (0.1% formic acid (FA) in water/acetonitrile (2:8, v/v)) after 42 min, 55% MS solvent B after 58 min and 99% MS solvent B at 60 min, followed by a 10-minutes wash at 99% MS solvent B and re-equilibration with MS solvent A (0.1% FA in water). The first 15 minutes, the flow rate was set to 750 nl/min after which it was kept constant at 300 nl/min.

The mass spectrometer was operated in data-dependent mode, automatically switching between MS and MS/MS acquisition. Full-scan MS spectra (300-1500 m/z) were acquired in 3 s acquisition cycles at a resolution of 120,000 in the Orbitrap analyzer after accumulation to a target AGC value of 200,000 with a maximum injection time of 250 ms. The precursor ions were filtered for charge states (2-7 required), dynamic range (60 s; ± 10 ppm window) and intensity (minimal intensity of 5E3). The precursor ions were selected in the ion routing multipole with an isolation window of 1.6 Da and accumulated to an AGC target of 10E3 or a maximum injection time of 40 ms and activated using CID fragmentation (35% NCE). The fragments were analyzed in the Ion Trap Analyzer at rapid scan rate.

The generated MS/MS peak lists were searched with Mascot using the Mascot Daemon interface (version 2.6.0, Matrix Science, Boston, MA). MS data were searched twice, once being matched against a custom-build database (containing UniProt, UniProt isoform entries appended with Ribo-seq derived protein sequences, generation and composition is described in detail in [3]) and once being matched against the UniProt database (release 2019_04). The Mascot search parameters were for both searches the same and as follows: heavy acetylation of lysine side-chains (with ^13^C_1_D_3_-acetate), carbamidomethylation of cysteine and methionine oxidation to methionine-sulfoxide were set as fixed modifications. Variable modifications were acetylation of N-termini (both light and heavy due to the ^13^C_1_D_3_ label) and pyroglutamate formation of N-terminal glutamine (both at the peptide level). The enzyme settings were: endoproteinase semi-Arg-C/P (semi-Arg-C specificity with Arg-Pro cleavage allowed) allowing for two missed cleavages for the trypsin sample. For chymotrypsin and GluC, the enzyme settings were semi-Chymo and semi-GluC. For GluC, two missed cleavages were allowed, while for chymotrypsin four missed cleavages were allowed. Mass tolerance was set to 10 ppm on the precursor ion and to 0.5 Da on fragment ions. In addition, the C13 setting of Mascot was set to 1. Peptide charge was set to 1+, 2+, 3+ and instrument setting was put to ESI-TRAP. Raw DAT-result files of MASCOT were further queried using ms_lims [4]. Only peptides that were ranked first and scored above the threshold score set at 99% confidence were withheld. The FDR was estimated by searching a decoy database (a reversed version of the custom generated database or the UniProt database), which resulted in an FDR of 0.44% for the trypsin sample, 0.14% for the chymotrypsin sample and 0.53% for the GluC sample for the custom-build database and FDRs of 0.72%, 0.29% and 1.13% for the trypsin, chymotrypsin and GluC sample respectively with the UniProt database.

*LC-MS/MS analysis of Virotrap samples and protein identification*

For each sample, 7.5 µl of the supernatant was injected for LC-MS/MS analysis on an Ultimate 3000 RSLCnano system in-line connected to a Q Exactive HF Biopharma mass spectrometer (Thermo Scientific, Germany). Trapping was performed at 10 μl/min for 4 min in loading solvent A on a 20 mm trapping column (made in-house, 100 μm internal diameter (I.D.), 5 μm beads, C18 Reprosil-HD, Dr. Maisch, Germany). The peptides were separated on a 250 mm Waters nanoEase M/Z HSS T3 Column, 100 Å, 1.8 µm, 75 µm inner diameter (Waters Corporation, UK) kept at a constant temperature of 50 °C. Peptides were eluted by a non-linear gradient starting at 1% MS solvent B reaching 55% MS solvent B in 80 min, 97% MS solvent B in 90 min followed by a 5-minute wash at 97% MS solvent B and re-equilibration with MS solvent A. The mass spectrometer was operated in data-dependent mode, automatically switching between MS and MS/MS acquisition for the 12 most abundant ion peaks per MS spectrum. Full-scan MS spectra (375-1500 m/z) were acquired at a resolution of 60,000 in the Orbitrap analyzer after accumulation to a target value of 3,000,000. The 12 most intense ions above a threshold value of 13,000 were isolated with a width of 1.5 m/z for fragmentation at a normalized collision energy of 30% after filling the trap at a target value of 100,000 for maximum 80 ms. MS/MS spectra (200-2,000 m/z) were acquired at a resolution of 15,000 in the Orbitrap analyzer.

The generated MS/MS spectra were processed with MaxQuant (version 1.6.17.0) using the Andromeda search engine with default search settings, including a false discovery rate set at 1% on both the peptide and protein level. For the Virotrap experiment, the sequences of the human proteins in the Swiss-Prot database (release Jan 2021) were complemented with the sequences of GAG, VSV-G, FLAG-VSVG, eDHFR and the sequences of the baits were replaced by the sequences of their shortest proteoforms (only the baits used in that set were adjusted) and used as search space. The replacement of the bait sequences to the shortest variant was done to allow more correct quantification of these proteins over the full-length and proteoform bait interactomes.

The enzyme specificity was set at trypsin/P, allowing for two missed cleavages. Variable modifications were set to oxidation of methionine residues and N-terminal protein acetylation. Other settings were kept as standard unless specified here. Proteins were quantified by the MaxLFQ algorithm integrated in the MaxQuant software [5]. Peptides with a minimum of two ratio counts of both unique and razor peptides were considered for protein quantification. Further data analysis was performed with the Perseus software [6] (version 1.6.15.0) after uploading the protein groups file from MaxQuant. Proteins only identified by site and reverse database hits were removed as well as potential contaminants. Replicate samples were grouped and proteins with less than three valid values in at least one group were removed. Missing values were imputed using imputeLCMD (R package implemented in Perseus) with a truncated distribution with parameters estimated using quantile regression (QRILC).

*Generation of clones for AP-MS*

For EIF4A1, MAVS and PAIP1, both the canonical proteins and N-terminal proteoforms pMET7-Bait-tag proteins were constructed via an in-house developed Golden Gate assembly platform.

The following table gives an overview of all generated constructs: pMET7_MAVS_FL_FLAG, pMET7_MAVS_PR_FLAG, pMET7_PAIP1_FL_FLAG, pMET7_PAIP1_PR_FLAG, pMET7_EIF4A1_FL_FLAG, pMET7_EIF4A1_PR_FLAG and pMET7_eDHFR-FLAG

1.	van Loo, G., et al., *Endonuclease G: a mitochondrial protein released in apoptosis and involved in caspase-independent DNA degradation.* Cell Death Differ, 2001. **8**(12): p. 1136-42.

2.	Staes, A., et al., *Protease Substrate Profiling by N-Terminal COFRADIC.* Methods Mol Biol, 2017. **1574**: p. 51-76.

3.	Bogaert, A., et al., *Limited Evidence for Protein Products of Noncoding Transcripts in the HEK293T Cellular Cytosol.* Mol Cell Proteomics, 2022. **21**(8): p. 100264.

4.	Helsens, K., et al., *ms_lims, a simple yet powerful open source laboratory information management system for MS-driven proteomics.* Proteomics, 2010. **10**(6): p. 1261-4.

5.	Cox, J., et al., *Accurate proteome-wide label-free quantification by delayed normalization and maximal peptide ratio extraction, termed MaxLFQ.* Mol Cell Proteomics, 2014. **13**(9): p. 2513-26.

6.	Tyanova, S., et al., *The Perseus computational platform for comprehensive analysis of (prote)omics data.* Nat Methods, 2016. **13**(9): p. 731-40.
